# Color-biased regions in the ventral visual pathway are food-selective

**DOI:** 10.1101/2022.05.25.493425

**Authors:** Ian Morgan Leo Pennock, Chris Racey, Emily Allen, Yihan Wu, Thomas Naselaris, Kendrick Kay, Anna Franklin, Jenny Bosten

**Author notes:** **Corresponding Author**: Ian Morgan Leo Pennock, The Sussex Colour Group, School of Psychology, University of Sussex, Brighton, United Kingdom.

## Abstract

The ventral visual pathway is well known to be involved in recognizing and categorizing objects (Kanwisher and Dilks, 2013). Three color-biased areas have also been found between face and place selective areas in the ventral visual pathway (Lafer-Sousa et al., 2016). To understand the function of these color-biased areas in a region known for object recognition, we analyzed the Natural Scenes Dataset (NSD; Allen et al., 2022), a large 7T fMRI dataset from 8 participants who viewed up to 30,000 trials of images of colored natural scenes. In a whole-brain analysis, we correlated the average color saturation of the images and the voxel responses, revealing color-biased areas that diverge into two streams in the ventral visual pathway, beginning in V4 and extending medially and laterally of the Fusiform Face Area in both hemispheres. We drew regions of interest (ROIs) for the two streams and found that the images for each ROI that evoked the largest responses had certain characteristics: They contained food, contained circular objects, had higher color saturation, contained warmer hues, and had more luminance entropy. A multiple linear regression showed that the presence of food in images was the strongest predictor of voxel responses in the medial and lateral color-biased regions for all eight participants, but that color saturation also contributed independently to voxel responses. Our results show that these areas are food-selective and color biased. We suggest that these streams might be involved in using color to recognize and judge the properties of food.

## INTRODUCTION

The ventral visual pathway is specialized for the perception and recognition of visual objects, e.g. faces (Kanwisher et al., 1997; Kanwisher and Yovel, 2006), places (Epstein et al., 1999; Epstein and Kanwisher, 1998), bodies (Downing et al., 2001; Peelen and Downing, 2007), and words (Dehaene and Cohen, 2011; Kay and Yeatman, 2017). Color is an important feature of objects (Gegenfurtner and Rieger, 2000; Witzel and Gegenfurtner, 2018), but its representation in the ventral visual pathway is not well understood.

The processing of color information begins in the retina with a comparison of the activities of the three classes of cone sensitive at short (S), medium (M) and long (L) wavelengths of light. Subsequently, different classes of retinal ganglion cells send luminance and color information to the lateral geniculate nucleus which projects to V1 (Conway et al., 2018). In the early visual cortices such as V1, V2, V3 and V4v color responsiveness has been studied using fMRI (Bannert and Bartels, 2018, 2013; Beauchamp et al., 1999; Brouwer and Heeger, 2009; Hadjikhani et al., 1998). V1 to V3 respond to color among other features (Mullen et al., 2007) while V4 and the ventral occipital region (VO; anterior to V4) are thought to be specialized for processing color (Mullen, 2019). Voxel activity patterns in V4, VO1 and VO2 can strongly distinguish chromatic from achromatic stimuli (Goddard and Mullen, 2020), and representational similarity analysis has provided evidence for a perceptual representation of color in these areas. More cognitive color tasks are also associated with V4, such as mental imagery for color (Bannert and Bartels, 2018) and color memory (Bannert and Bartels, 2013). As color information progresses through visual cortical regions, its representation likely becomes transformed to aid cognitive tasks such as object perception (Vandenbroucke et al., 2014).

Most studies of color perception present simple stimuli such as color patches, rather than color as it naturally occurs, embedded in natural scenes. However, in daily life our visual system encounters colors as part of a conjunction of object features integrated in context within natural scenes. With simple stimuli color is dissociated from its regular context and meaning: These stimuli have basic spatial form, may be selected from a restricted color gamut, and are typically presented on a uniform surround. The visual responses to carefully controlled colored stimuli might be quite different to those that occur in response to colors in their complex, naturalistic settings. For example, for colored patches, decoding accuracy drops between V1 to V4 (Bannert and Bartels, 2018; Brouwer and Heeger, 2009), while for colored object categories decoding accuracy increases through the same areas (Vandenbroucke et al., 2014). To understand how the brain represents color in its usual context, it is therefore crucial to use complex stimuli such as natural scenes.

Only one existing study has addressed human neural color representation using complex stimuli. Lafer-Sousa et al. (2016) presented colored and greyscale videos of faces, bodies, places, objects, and scrambled scenes and found posterior, central and anterior color-biased regions located between place- (parahippocampal place area; PPA) and face- (fusiform face area; FFA and occipital face area; OFA) selective areas. One interpretation of these color-biased regions is that they are specialized for color irrespective of object category and spatial form. However, they are located in the ventral visual pathway which is known to be responsive to a range of complex visual objects (Downing et al., 2001; Epstein et al., 1999; Kanwisher et al., 1997; Kanwisher and Yovel, 2006; Kay and Yeatman, 2017; Lafer-Sousa et al., 2016; Peelen and Downing, 2007). It is therefore possible that the color biases observed in these regions are attributable to the color properties of particular preferred object classes.

We aimed to characterize the neural representation of color in the context of objects in natural scenes. The Natural Scenes Dataset (NSD; Allen et al., 2022) provides a unique opportunity for this endeavor. It is an unprecedented large-scale fMRI dataset in which participants viewed thousands of colored (and some greyscale) natural scenes over 30 to 40 sessions in a 7T fMRI scanner. This dataset therefore has impressively high signal-to-noise which enables excellent statistical power (Naselaris et al., 2021). Images of natural scenes are highly dimensional and visual features correlate strongly, which makes the contributions of different features difficult to disentangle. With its huge number of well-characterized and segmented stimulus images, the NSD is one of the best datasets currently available to uncover the neural representations underlying perception of natural scenes (Allen et al., 2022; Lin et al., 2014). We found two streams in the ventral visual pathway that showed responses to the color properties of the NSD images. We found that both streams were primarily responsive to food objects, implying that color is a key part of the neural representation of food in these ventral visual areas.

## RESULTS

### Correlation with saturation

We conducted a whole brain correlation between the average color saturation of each NSD image and the percentage BOLD signal change (Figure 1), to locate brain areas which respond to color. Since saturation and luminance (Figure 2A and SI Figure 1) are correlated in natural scenes (Long et al., 2006), we used average luminance (quantified as L+M) of the pixel values, without a dark filter applied, as a covariate. The correlations were Bonferroni corrected for each participant based on the number of voxels in participant-native space. We also conducted an analysis separately for odd and even images to measure split-half reliability.

**Figure 1:**
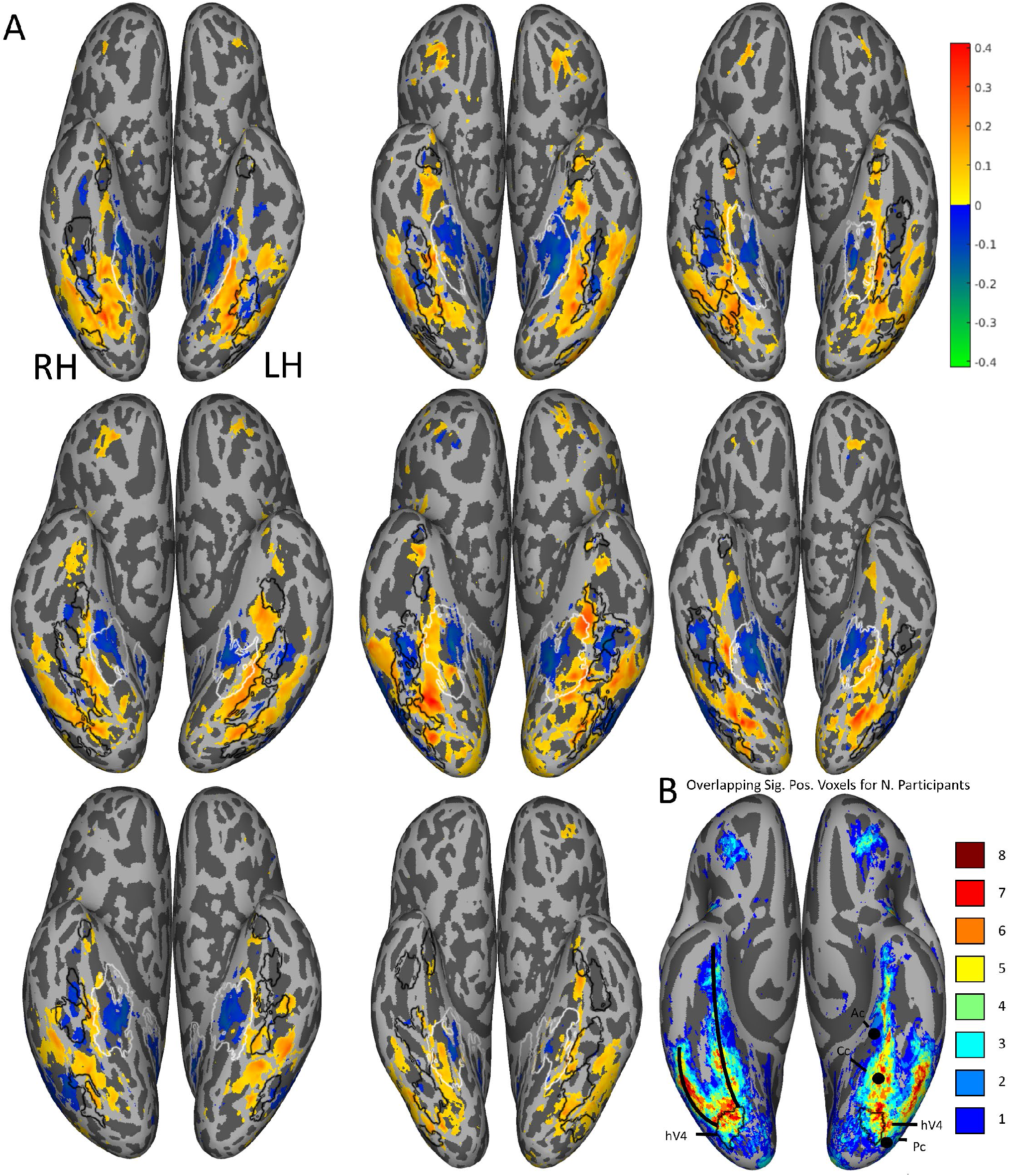
Correlation between average saturation and voxel responses. (A). Eight (top-left is participant 1, bottom-middle is participant 8; going left to right) Pearson correlation maps of the ventral view in native participant-space. The maps show for each voxel the correlation between the mean saturation of each image and the corresponding brain responses, with mean luminance as a covariate. Positive correlations are displayed in red and yellow, and negative correlations in green and blue. Only correlations with significant Bonferroni-corrected p-values are shown. Black contours indicate face-selective brain regions for each individual participant (FFA-1, FFA-2, OFA, mTL-faces and ATL-faces) and white contours indicate place-selective areas for each individual participant (PPA and RSC); for a description of how these regions were defined see Allen et al. (2022). (B). The number of participants showing overlapping significant positive voxels in fsaverage space. On the right hemisphere, the medial and lateral ROIs in the ventral visual pathway are indicated (solid black lines) and on the left hemisphere the coordinates of the color-biased regions identified by Lafer-Sousa et al. (2016) are shown (Ac, Cc, Pc). For both hemispheres hV4 from the brain atlas by Wang et al. 2015 are indicated by the black contours.

**Figure 2:**
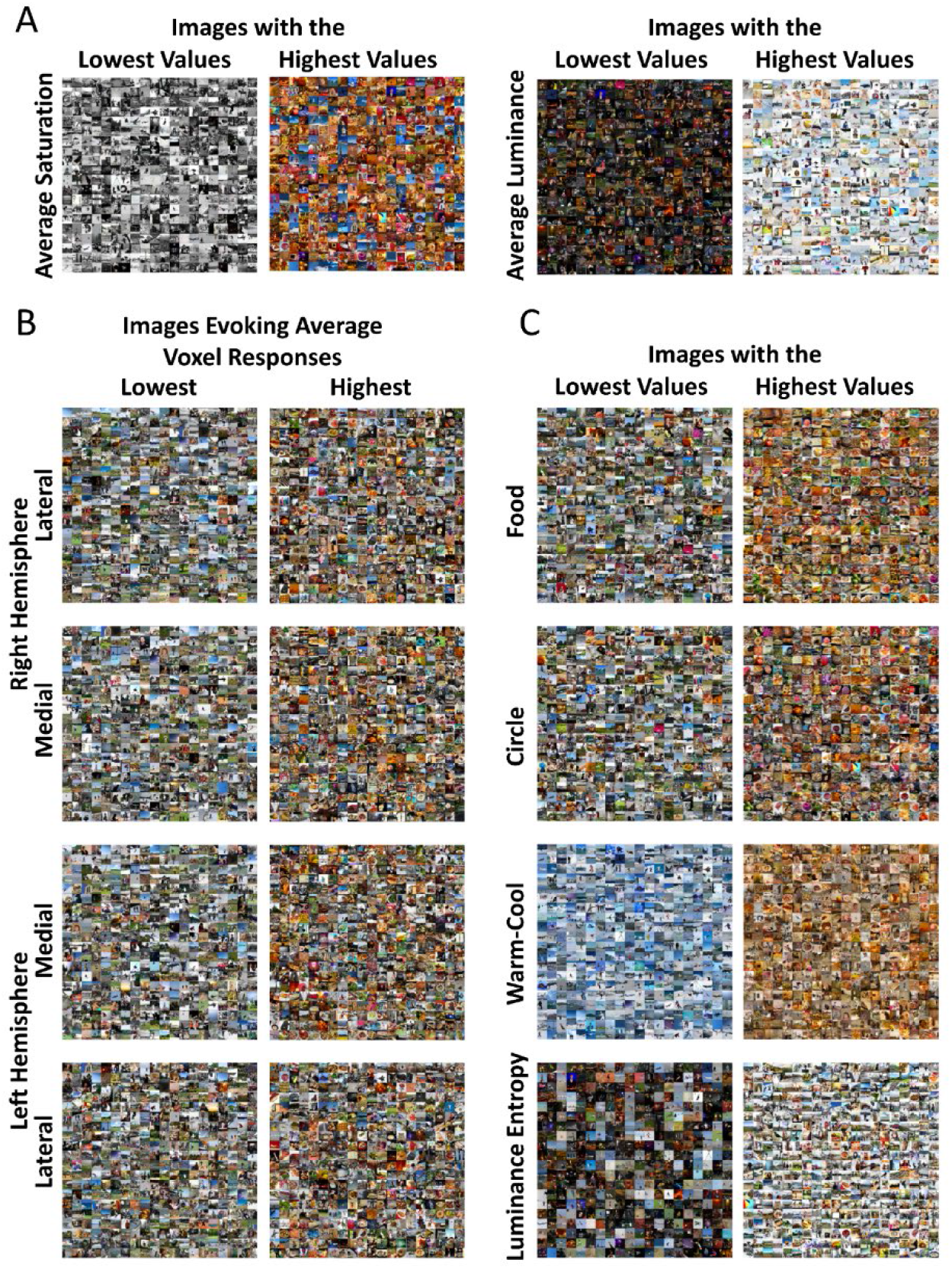
Montages of each image statistic and images evoking the highest and lowest ROI responses. Montages of the 400 images with lowest (left) and highest (right) of participant 1. (A) Montages of the image statistics included in the correlation analysis: average saturation (based on the NSD grey background. Each pixel saturation level was measured comparing the distance between the values in Macleod-Boynton chromaticity diagram and the NSD grey background) and average luminance. (B) Montages show the images for the lateral and medial ROI for both hemispheres that have the highest and lowest averaged z-scored voxel responses for participant one. The correlation maps were used to draw Regions of Interest and then the selected voxels were averaged for each image. The 400 images evoking the highest and lowest voxel responses were selected for each montage (from top to bottom: right hemisphere-lateral ROI, right hemisphere-medial ROI, left hemisphere-medial ROI, left hemisphere-lateral ROI, and the left row is the lowest responses and, the right row is the highest responses. (C) For the multiple linear regression four image statistics were added (together with average saturation and average luminance). Montages of images statistics that were added to the multiple linear regression: food, circle, warm-cool ratings and luminance entropy in the NSD dataset. The left row is the lowest responses, and the right row is the highest responses.

For all participants there were areas showing positive correlations between saturation and voxel responses in the ventral visual pathway, beginning in V4, and diverging into two distinct medial and lateral regions of interest (ROIs; Figure 1). The medial ROI is located between face and place areas (Figure 1; see fLoc-experiment by Allen et al., (2022) for the category-selective regions) and is roughly in agreement with the location of the color-biased regions identified by Lafer-Sousa et al. (2016) (Figure 1B). The split-half analysis showed high reliability, with r = 0.82 (range = 0.71 – 0.89 for different participants) when voxel activities were correlated over the whole brain.

For all 8 participants there were also areas that showed negative correlations between saturation and voxel responses, in the PPA (Figure 1A) and for the region located between the lateral and medial ROIs that showed positive correlations (Figure 1A). For seven participants there was an area of negative correlation lateral of the lateral ROI, roughly corresponding to area MT. For six participants (and one further participant in the left hemisphere only) there was a positive correlation with saturation in prefrontal regions (Figure 1A).

### Montages of images producing the highest and lowest voxel responses

Our correlation analysis between BOLD and saturation revealed areas responsive to color in the ventral visual pathway for all participants. To better understand how the image features these areas respond to we created montages of the images that evoked the highest and lowest voxel responses for these areas, split into four ROIs (medial and lateral, left and right hemispheres; Figure 2B for participant 1 and SI figure 2 for other participants). For a description of how the ROIs were defined, see Methods.

By inspecting the montages, we identified multiple image properties present in images evoking the highest responses but not in images evoking the lowest responses. These properties were food such as bananas, donuts, and pizzas; circular objects such as plates, clocks and stop signs; warm colors such as reds and oranges; and luminance entropy (how well luminance values in one location can predict the values in nearby locations; Mather, 2020). These image characteristics were consistent across all participants, medial and lateral ROIs, and hemispheres, suggesting that the four ROIs all process a similar type of visual information.

### Image statistics and their intercorrelations

We calculated image statistics for the image properties that appear to distinguish the images that evoke the higher and lower voxel responses in our ROIs. We also included average luminance in the intercorrelation analysis as it was used as a covariate in the correlation analysis for saturation. Our image statistics were pixel count for food objects, pixel count for circular objects, warm-cool ratings, average saturation, luminance entropy and average luminance (see Methods for a detailed description).

The six image statistics were significantly intercorrelated (see Supplementary Figure 4 for correlation matrices for each participant and Figure 2A and C for montages). Average luminance and luminance entropy were strongly positively correlated (group average ρ = 0.68), and circular objects and food images were moderately correlated (group average ρ = 0.37). All but one other pairs of image statistics had low but significant correlations (group average ρ < 0.30). Circular object pixel counts and luminance entropy were not significantly correlated for seven of the eight participants. The relationships between image statistics were highly consistent between participants who viewed different image sets (0.9991 ≤ ρ ≤ 0.9999 for pairwise correlations between image statistic correlation matrices between participants).

### Image statistics and average ROI responses

To investigate the relationship between each image statistic and average voxel responses for our four ROIs (medial and lateral areas in both hemispheres), we plotted moving average ROI responses against each the image statistic (Figure 3A). ROI responses show positive linear relationships with average saturation and warm-cool ratings. ROI responses show a positive non-linear (decelerating) relationship with food pixel count and circular object pixel count, with a higher gain for food pixel count than for any of the other image statistics. There is no relationship between ROI responses and luminance entropy, and a small negative relationship between ROI responses and average luminance. These findings are consistent across hemispheres and ROIs for all eight participants (see SI Figure 3 for results for individual participants).

**Figure 3:**
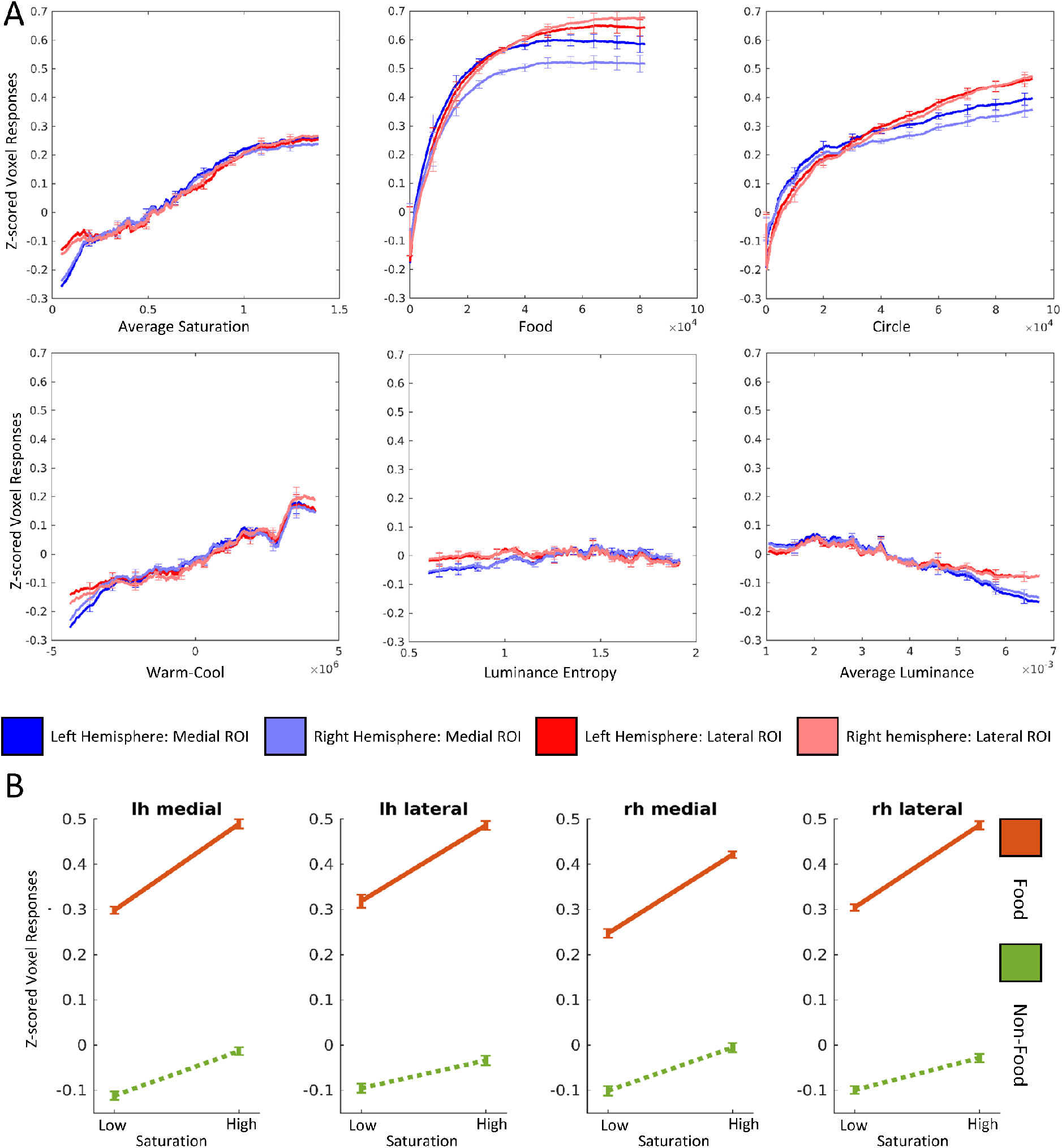
Average ROI responses for the image statistics and two-way ANOVA. (A) The average z-scored voxel response in the medial and lateral ROI of the left and right hemisphere. The x-axis shows the z-scored averaged voxel response, and the y-axis shows an image statistic: average saturation per image (distance from the NSD grey background in MacLeod-Boynton space), number of pixels that contain an image of food, number of pixels that contain an image of a circular object, summed warm-cool ratings based on individual pixel ratings, luminance (L+M) entropy (Mather, 2020), average luminance values for each image. The image statistics were sorted from lowest to highest trials based on the image statistic. Then the average of 500 z-scored voxel responses were sequentially presented: 1-500, 2-501, 3-502. We interpolated the data and then averaged across all participants. Each participant saw 8,302-9000 unique images and a direct comparison without interpolation could not be made. Error bars are the 95% confidence intervals within participants. The un-interpolated version of the individual participants can be found in SI figure 3. (B) A two-way ANOVA with food and saturation as factors was conducted on z-scored voxel response for all four ROIs (left (lh) and right (rh) hemisphere, and medial and lateral ROI). On the x-axis, the low and high saturation image groups are displayed. The y-axis displays the z-scored average voxel response. The orange line represents images that contained food and the green line are images that do not contain images of food based on the COCO categories. Error bars are the 95% confidence intervals within participants.

### Multiple linear regressions

We included the six image statistics in a multiple linear regression to identify the best predictors for the average voxel responses for our four ROIs. A regression analysis showed significant relationships in all four ROIs (Medial ROI LH: mean F (6,9648) over 8 participants = 283, SD = 99.7, p < 5.43 × 10^−188^, mean R^2^ = 0.15, SD = 0.04; Medial ROI LH: mean F (6,9648) = 239, SD = 100.0, p < 3.09 × 10^−169^, mean R^2^ = 0.13, SD = 0.04; Lateral ROI LH: mean F (6,9648) = 218, SD = 94.8, p < 5.60 × 10^−115^, mean R^2^ = 0.12, SD = 0.04; Lateral ROI RH: mean F (6,9648) = 210, SD =96.6, p < 3.19 × 10^−122^, mean R^2^ = 0.11, SD = 0.04). Summary results in Table 1 show that food is the strongest predictor for all four ROIs in all eight participants. Individual results for each participant are available in SI Table 2.

**Table 1.** Multiple Linear Regression Beta coefficients.

|  | B0 | Average Saturation | Food | Circles | Warm-Cool | Luminance Entropy | Average Luminance |
| --- | --- | --- | --- | --- | --- | --- | --- |
| Medial Left | $5.3863 \times 10^{-9}$ | 0.0577 | 0.2060 | 0.0479 | 0.0337 | 0.0440 | - 0.0511 |
| Medial Right | $5.4712 \times 10^{-9}$ | 0.0568 | 0.1702 | 0.0474 | 0.0310 | 0.0433 | - 0.0468 |
| Lateral Left | $9.9330 \times 10^{-12}$ | 0.0350 | 0.2384 | 0.0364 | 0.0288 | - 0.0053 | - 0.0022 |
| Lateral Right | $1.7063 \times 10^{-9}$ | 0.0397 | 0.2408 | 0.0233 | 0.0450 | - 0.0093 | 0.0014 |

The average beta coefficients for each image statistic in the multiple linear regressions with average voxel responses for each of the four ROIs.

### Multiple linear regressions on individual voxels

We also ran the multiple linear regression on all the voxels that showed a significant positive correlation with saturation (all voxels were significant, *p < 0.01*). For each voxel we ranked the highest image statistic based on its beta coefficient. Figure 4 shows the first ranked imaged statistic for each voxel. Food pixel count produced the first ranked beta coefficient in the single-voxel multiple regressions for almost all voxels, suggesting that food is the strongest predictor for all four ROIs even at an individual voxel level. For the left medial, left lateral, right medial and right lateral ROIs respectively, 81%, 93%, 73% and 93% of voxels had food as the strongest predictor, and only 4%, 0.6%, 6% and 2% had saturation as the strongest predictor. For the other image statistics there was no consistent pattern. Voxels activity in early visual areas was strongly predicted by luminance entropy. For V1 voxels defined by the HCPMMP 1.0 atlas (Glasser et al., 2016) 80% of voxels in V1 that had a significantly positive relationship with luminance entropy for the first ranked beta coefficient, 9% had food and 3% had saturation.

**Figure 4.**
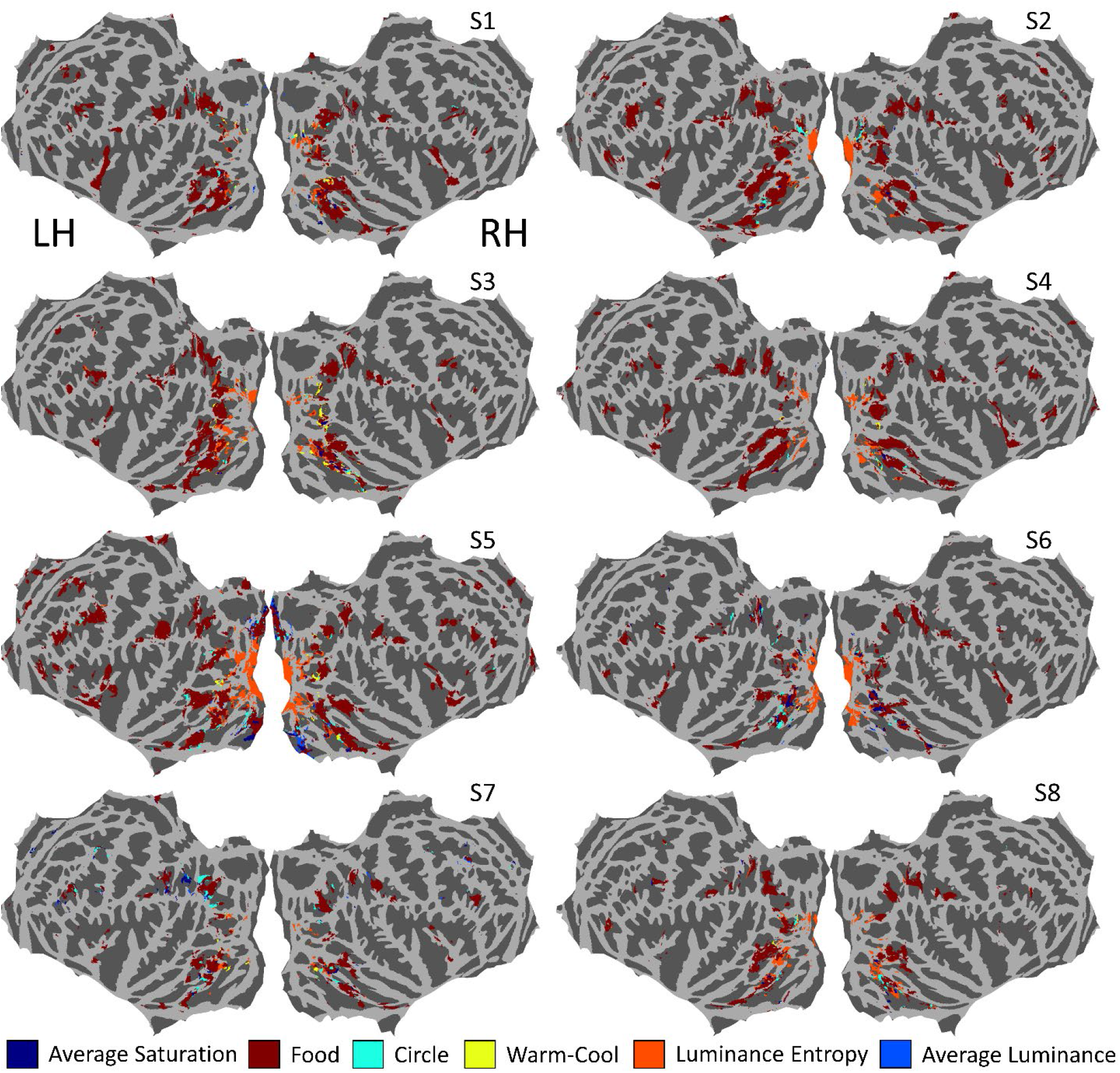
Multiple linear regression on individual voxels. For each of the eight participants are shown flattened whole-brain cortical maps in fsaverage space, in which each voxel that showed a significantly positive correlation with saturation is colored. For each voxel, the color is according to the image statistic that had the largest beta coefficient in the multiple regression for that voxel.

### ANOVA with saturation and food

The multiple linear regressions for the ROIs showed that food pixel count had the highest beta coefficients of the six image statistics. The results of our whole-brain correlation with saturation and previous literature (Lafer-Sousa et al., 2016) imply that these area are responsive to color. We therefore sought to further investigate the contributions of saturation and food to ROI responses by conducting a two-way ANOVA (Figure 3B). For all four ROIs the ANOVA revealed a significant main effect for food for all 8 participants (medial area LH: mean F (1, 9651) = 596, p < 1 × 10^−323^; medial area RH: mean F (1, 9651) = 464, p < 1 × 10^−323^; lateral area LH: mean F (1, 9651) = 540, p < 1 × 10^−323^; lateral area RH: mean F (1, 9651) = 493, p < 1 × 10^−323^). All four ROIs also showed a significant main effect of saturation for all 8 participants (medial are LH: mean F (1, 9651) = 60.1, p < 1.78 × 10^−8^; medial area RH: mean F (1, 9651) = 55.2, p < 4.01 × 10^−8^; lateral area LH: mean F (1, 9651) = 32.5, p < 1.73 × 10^−4^; lateral area RH: mean F (1, 9651) < 34.9, p < 5.41 × 10^−7^). There were significant interactions for some participants in some ROIs (5 for the medial area in the LH, 4 for the medial area in the RH, 3 for the lateral area in the LH, and 6 for the lateral area in the RH). For ANOVA results for all participants, see SI Figure 5 and SI Table 1. For a heatmap between food and saturation see SI Figure 6.

### Food versus non-food

Our results showed food images to be a strong predictor of responses in our ROIs, but since they were defined by responses to saturation rather than food, the analyses reported so far could miss voxels that respond to food but not to saturation. We therefore conducted an analysis of the differences between responses to stimuli that contain images of food and responses to images that do not contain food. Each participant (1 to 8) saw 1284, 1284, 1176, 1237, 1303, 1240, 1309, 1127 images of food respectively. All other images were considered non-food images based on the Microsoft’s Common Objects in Context (COCO; Lin et al., 2014) food categories. Figure 5 shows results plotted for the whole brain, also including the peak activation coordinates of a fMRI meta-analysis of food images (van der Laan et al., 2011) in the right hemisphere. We converted the Bonferroni-corrected threshold for the saturation correlation analysis (Figure 1) to T-values and applied the same threshold to Figure 5 to make a comparison possible.

**Figure 5.**
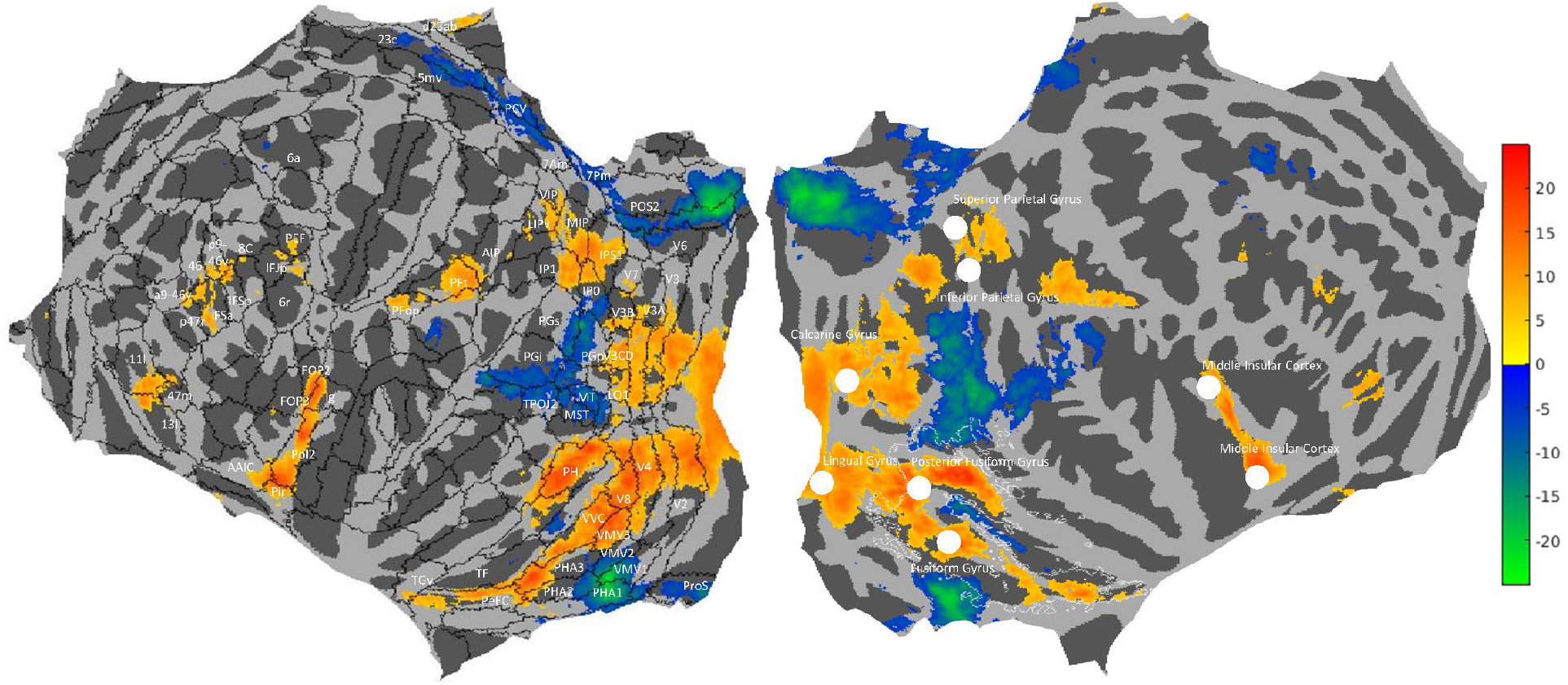
Analysis of responses to food versus non-food images. A flattened cortical map in fsaverage space showing t-statistics for the differences between average voxel responses for food versus non-food images. The Human Connectome Project Atlas (HCP_MMP1; Glasser et al., 2016) is overlayed for left hemisphere (black contours), with regions labelled where they contained voxels with significant t-statistics. On the right hemisphere are plotted coordinates identified by van der Laan et al. (2011) in a meta-analysis of brain areas responsive to food indicated by a white dot (see their Table 2) and contours of the medial and lateral ROIs of Figure 1B in white.

Our results show that food images significantly activate similar areas to the ROIs we identified for their correlated activity with saturation (see white contours superimposed on the RH in Figure 5). The Activation Likelihood Estimation (ALE) meta-analysis by van der Laan et al., (2011) identified locations in the Fusiform Gyrus and Posterior Fusiform Gyrus that are responsive to food, which are located in the medial and lateral ROIs. According to the Human Connectome Project atlas (HCP-MMP 1.0 atlas; Glasser et al., 2016; see black contours superimposed on the LH in Figure 5), the medial ROI ends in the perirhinal ectorhinal cortex (PeEC) and the lateral ROI ends in area Ph.

There is also activation in the early visual areas (V1, V2, V3 and V4) which is unlikely to be driven by food itself but by luminance entropy (Figure 4) correlated with the presence of food in the NSD stimulus set. There is activation in dorsal areas of the visual cortex (V1, V2, V3 and V4) to V3CD, LO1 and V3B. Another cluster of activation is found in IPS1, IP1 and IP0 and MIP, VIP, LIPv, the latter cluster was also identified in the ALE meta-analysis. Two more areas of activation are found in PFt and PFop and part of AIP in both hemispheres, which the ALE meta analysis identified in the left hemisphere only (Inferior Parietal Gyrus). Another area of activation is found in Pol2, Ig, MI, AAIC, Pir, FOP2 for both hemispheres and a part of FOP3 for the left hemisphere, which corresponds to the Insular cortex in both hemispheres. Some smaller clusters of activation are found in PEF in both hemispheres, in the left hemisphere also spanning parts of IFJp and 6r. Responses to non-food images are significantly higher than to food images in areas MT, MST, TPOJ1, TPOJ2, TPOJ3, PGi, PGp, PGs, IP0, STV, PSL, and PF which cluster together. This is also the case in POS1, POS2, DV, PCV, 5mv, 23c, and another in VMV1, PH1 and ProS.

### ROI responses to object categories

We investigated the average voxel responses for each ROI to each object category in the COCO dataset. We found that on average all food objects provoke the highest voxel responses (Figure 6 and SI Figure 7 for individual participants). Objects involved in food preparation and consumption such as spoons, knives, forks and dining tables also provoked high voxel responses. Results suggest a variety of food objects and food-associated objects drive our results.

**Figure 6.**
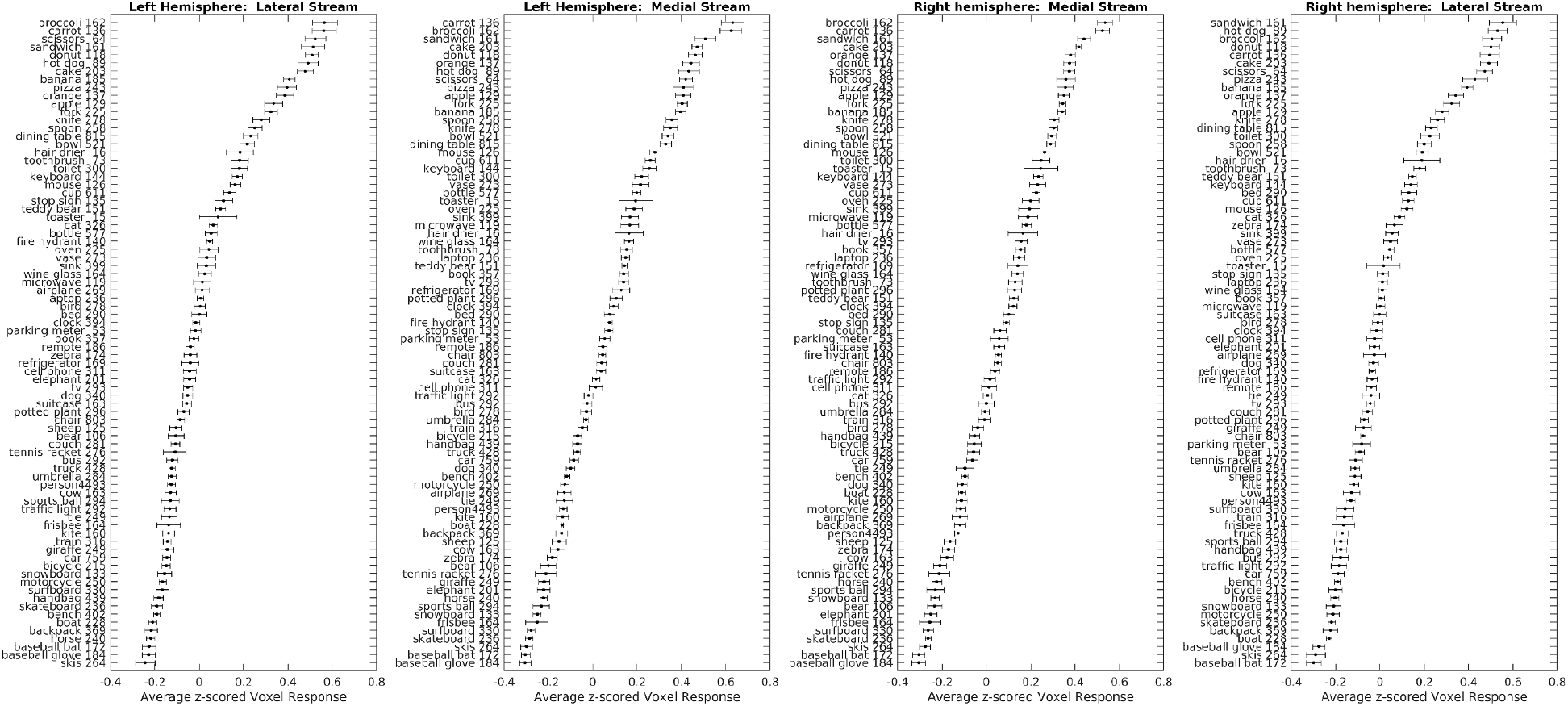
Average voxel responses in the four ROIs for each COCO object category. The four panels show average voxel response to each object category, with one panel for each ROI. The x-axis displays the average Z-scored voxel response and the y-axis shows the object name and the average number of images across 8 participants that contained this object, rounded. The 80 object categories are those identified and segmented in the COCO dataset. The object categories are ordered from those provoking the strongest response (top) to those provoking the weakest response (bottom). There may be multiple objects in one image. Error bars are between-subject standard errors of the mean.

## DISCUSSION

Our results show that color-biased regions in the ventral visual pathway are food-selective. We identified four ROIs in the ventral visual pathway responsive to the average color saturation of images, one located medially and the other laterally of the FFA in each hemisphere. The NSD dataset enabled an in-depth analysis of the responsiveness of these color-biased regions because of the large number and variety of images of natural scenes presented in the scanner. When we investigated a variety of image characteristics we found the color-biased areas to be most strongly activated by food, with a smaller independent response to saturation. Our results lead us to reinterpret the regions as food-selective but color-biased, implying that color is important in the neural representation of food. Our results uncover a visual stream for food and food associated objects in the ventral visual pathway not previously identified in the visual neuroscience literature, although some regions within this stream have been identified in a meta-analysis on fMRI studies of food (van der Laan et al., 2011). The ventral visual pathway is known to contain sub-streams for processing faces, places, bodies, and words: our results suggest a prominent role for food selectivity as well.

### Reliability and consistency of results

We conducted a split half reliability analysis over odd and even images for our correlation between saturation and voxel responses which showed strong reliability over the whole brain (mean r = 0.82, range = 0.71 – 0.89 for different participants). The montages of the images evoking the highest responses in our ROIs contained similar image features for all eight participants (Figure 2B and SI Figure 2), which were absent in the montages of images that evoked the lowest responses. The intercorrelations between image statistics were similar for all participants, suggesting that there were no major differences between the unique images shown to each participant to take into account when interpreting the results. Multiple analyses: plots of the relationships between image statistics and voxel responses (Figure 3A), and the multiple linear regresions for the ROIs (Table 1) and for the individual voxels (Figure 4) all showed that food was the strongest predictor of voxel responses in the ROIs. We therefore interpret these regions as food-selective.

### Color-biased regions in the ventral visual pathway

We found that the medial and lateral food-selective streams are still biased to color even in the absence of food. This is in line with the results of Lafer-Sousa et al. (2016), who showed no food stimuli in their fMRI experiment but found color-biased anterior, central, and posterior areas medial in the ventral visual pathway. Their findings also hinted at a lateral color-biased area for a few of their participants. Our results for all eight participants show two approximately continuous streams, diverging medially and laterally beginning in V4. We found that the medial stream extends further anteriorly from the anterior color-biased region identified by Lafer-Sousa et al. Our finding that the color-biased regions are selective for food, changes the existing interpretation of what these streams are functionally specialized for. Lafer-Sousa et al. proposed that the color-biased regions are specialized for color and are anatomically segregated from neighboring regions specialized for object processing, but we find instead that the color-biased areas are specialized for processing images of food. The smaller independent responses we find for these regions to saturation may have several different interpretations, which we discuss further below. In addition, we found negative correlations between saturation and the voxel responses (Figure 1A), mostly in areas that respond to faces, places, and motion; one possible explanation is that these image features tend to be associated with lower saturation.

### Food-Selective Streams in the Ventral Visual Pathway

Our results show lateral and medial food-selective streams in the ventral visual pathway. Previous studies have also found responsiveness to food in these areas, summarized in a meta-analysis by van der Laan et al. (2011). Although the meta-analysis of visual processing of food images shows an area of concordance between studies in the fusiform gyrus, some studies have looked for food-selective areas in the ventral visual pathway but did not find them (Downing et al., 2006; Pinsk et al., 2009), and Adamson and Troiani (2018) found that the left fusiform cortex responds equally well to faces and food. Our results imply that substantial neural resources in the ventral visual pathway are allocated to the important task of visually processing food.

Numerous studies have demonstrated a distinction between the processing of animate versus inanimate objects in the ventral visual pathway (Grill-Spector and Weiner, 2014; Martin et al., 1996), specifically that areas medial of the FFA respond preferentially to animate objects but lateral areas to inanimate objects (Grill-Spector and Weiner, 2014; Martin et al., 1996). At first glance, the existence of two food-selective streams separated by the FFA might appear to contradict this theory. However, the placement of food in the category distinction between animate and inanimate objects is ambiguous. For example, fruit and vegetables are living entities and foods, but pizzas and hot dogs are non-living foods processed from ingredients derived from living entities (Crutch and Warrington, 2003). However, when we calculated the average BOLD percentage signal change for images containing each object from each COCO category we found no clear distinction between the food categories (Figure 6). Further research is needed to understand the functional differences (or lack thereof) between the lateral and medial food-selective streams and whether they support a distinction between animate and inanimate objects.

The streams in the ventral visual pathway that we have identified as food-selective respond to all categories of food in the COCO image set (Figure 6), including fruits and vegetables as well as processed foods that were not available in the evolutionary past. We therefore speculate that the food-selective streams are tuned by exposure to food during a person’s lifetime. This would be analogous to within-lifetime tuning of the visual word form area, which, owing to the relatively recent development of written language, is unlikely to be innately specified (Baker et al., 2007; Kravitz et al., 2013). However, as the visual word form area is highly consistent across individuals, it also seems unlikely that it is formed solely through experience (Kravitz et al., 2013).

The results of our analysis of the responsiveness of the streams to different object categories (Figure 6) does not support a clear-cut distinction by the streams for the boundary between food and non-food objects, but instead supports more graded responses to food. This is suggested by the relatively high responses of the streams to food-associated objects such as spoons, forks, knives, and dining tables.

We must also consider the possibility that our results may be influenced by attention or expertise (Bilalic et al., 2016; Bukach et al., 2006; Gauthier et al., 1999; O’Craven et al., 1999; Xu, 2005). Participants may have been more attentive to images containing food than they were to images containing other objects. Figure 6 shows responses of the medial and lateral streams to images containing objects that could be considered attention-grabbing such as bears, baseball bats, and stop signs. However, these objects are not among those causing the greatest responses. Therefore, we consider it unlikely that these streams are driven by attention rather than food. Alternatively, food images may strongly activate these areas if they are general object processing areas but people have particular expertise for food. Again, we believe that the preponderance of food but also food-associated objects among those provoking the highest voxel responses renders this account unlikely.

### Food and Color in the Ventral Visual Pathway

We refer to the streams as ‘food-selective’ as they respond to food more strongly than they respond to color. However, our saturation findings suggest that they also respond to color. Why are these food-selective areas also color-biased? One possibility is that the medial and lateral streams may have two distinct specialisms unrelated to a common visual function, for both food and color. An alternative possibility, which we favor, is that these areas respond to collections of visual features common to food objects and also to these features even in the absence of explicit food objects, e.g., colors that are normally predictive of the presence of food. Humans use color as a heuristic for evaluating food (Foroni et al., 2016), and there is strong evidence that trichromatic vision helps animals to detect food (Osorio and Vorobyev, 1996; Regan et al., 2001; Sumner and Mollon, 2000a) and to judge its ripeness (Sumner and Mollon, 2000b). Visual information in the color domain is therefore central to detecting and recognizing food, so it seems plausible that food-selective regions would also respond to color as a relevant visual feature.

Conway (2018) proposed a tripartite organization of the ventral visual pathway, consisting of parallel streams for faces, places, and color, based on the results of studies in both humans and macaques (Lafer-Sousa et al., 2016; Lafer-Sousa and Conway, 2013; Verhoef et al., 2015). Humans share a common ancestor with macaques 25 million years ago (Conway, 2018) and as the human brain is a product of evolution, it is important to consider these systems from an evolutionary perspective as this might inform key principles of cortical organization (Cisek and Hayden, 2021). Our present findings for humans do not change the proposed organization of the ventral visual pathway but rather replace the idea of color specialization with specialization for food.

### Conclusion

The NSD is a one-of-a-kind dataset which allowed us to apply techniques that would not be informative for datasets with lower signal to noise. The high resolution and large number of trials provide strong evidence that color-biased regions in the ventral visual pathway are food-selective and that there are two distinct medial and lateral food-selective streams in both hemispheres which diverge from V4 and surround the FFA. Our results also suggest that these food-selective streams also respond to color but to a lesser degree. This finding redefines our understanding of color-biased regions in the ventral visual pathway and elaborates on its function and complexity in responding to both to food and to color.

## POSTSCRIPT

At the time of uploading on Biorxiv, we were made aware of a pre-print posted the day before by Jain et al. (2022) also using the NSD to explore food selectivity in the ventral visual pathway. Their findings about food selectivity are largely in agreement with our own. In distinction, our paper explores the relationship between color and food representation in the ventral visual pathway and shows that food-associated tools are also represented preferentially by the food-selective areas identified in both papers. Therefore, aside from food selectivity, our paper provides unique insight into color representation in the ventral visual pathway.

## MATERIALS AND METHODS

### NSD methods

Here we will provide an outline of the methods used to gather the NSD that are relevant for our analyses. Further detailed methods for the NSD can be found in Allen et al. (2022).

### Participants and MRI data acquisition

Eight participants were included in the study (six females; age range 19-32). All participants had normal or corrected-to-normal vision. Informed consent was obtained, and the University of Minnesota Institutional Review Board approved the experimental protocol. The participants were scanned using a 7T Siemens Magnetom passively shielded scanner at the University of Minnesota. A single channel transmit 32 channel receive RF head coil was used. The procedure gathered a gradient-echo EPI sequence at 1.8 mm isotropic resolution (whole brain; 84 axial slices, slice thickness 1.8mm, slice gap 0 mm, field-of-view 216 mm (FE) x 216 mm (PE), phase-encode direction anterior-to-posterior, matrix size 120 × 120, TR 1600 ms, TE 22.0 ms, flip angle 62°, echo spacing 0.66 ms, bandwidth 1736 Hz/pixel, partial Fourier 7/8, in-plane acceleration factor 2, and multiband slice acceleration factor 3).

### Stimulus presentation

A BOLDscreen 32 LCD monitor (Cambridge Research Systems, Rochester, UK) was positioned at the head of the scanner bed. The spatial resolution was 1920 pixels x 1080 pixels and the temporal resolution 120 Hz. The participants saw the monitor via a mirror mounted on the RF coil. There was a 5 cm distance between the participants’ eyes and the mirror and a 171.5 cm distance from the mirror to image of the monitor. A PR-655 spectroradiometer (PhotoResearch, Chatsworth, CA) was used to measure the spectral power distributions of the display primaries. The BOLDscreen was calibrated to behave as a linear display device which allowed us to calculate the transformation from RGB to LMS tristimulus cone activities. A gamma of 2 was applied to the natural scene images to approximate the viewing conditions of standard computer displays.

### Experimental task

The participants performed a long-term recognition task in which they had to press a button stating whether the scene presented on each trial had been shown before or not. On every trial a distinct image was shown for 3s with a semi-transparent red fixation dot (0.2° x 0.2°; 50% opacity) on a grey background (RGB: 127,127,127; S/(L+M) = 1.1154, L/(L+M) = 0.6852). After the 3s stimulus presentation the same fixation dot and the grey background were shown alone for 1s. Participants could respond any time during the 4s trial. Each run contained 75 trials (some of these were blank trials) and lasted 300s. There were twelve runs per session.

### Images displayed

73,000 distinct images were used which were a subsample (the train/val 2017 subsections) of COCO image dataset (Lin et al., 2014), which contains complex natural scenes with everyday objects in their usual contexts. The COCO dataset contains 80 object categories ranging from faces and cars, to food and stop signs (for examples see Figure 2). The images were 425 × 425 pixels x 3 RGB channels which were resized to fill 8.4 by 8.4 degrees on the BOLDscreen 32 display using linear interpolation. Participants had up to 40 scan sessions (range 30-40) and saw up to 10,000 images 3 times across these sessions: 8,302 – 9,000 of these were unique images and 907 – 1,000 images were seen by all participants.

### Preprocessing

The preprocessing of the functional data included temporal resampling, which corrected for slice time acquisition differences and upsampled the data to 1.000s. Field maps were acquired and the resampled volumes were undistorted using the field estimates. These volumes were used to estimate rigid-body motion parameters using SPM5 spm_realign. To correct for the head motion and spatial distortion, a single cubic interpolation was performed on the temporally resampled volumes. The mean fMRI volume was calculated and was corrected for gradient nonlinearities. Then the volume was co-registered to the gradient-corrected volume from the first scan session, so the first scan session was used as the target space for preparing fMRI data from the different scan sessions.

A GLM analysis was applied to the fMRI time-series data to estimate single-trial beta responses. The third beta version (b3, ‘betas_fithrf_GLMdenoise_RR’; native surface space) was used, and no alterations were made to this beta version’s preprocessing steps described in Allen et al. (2022). In brief, the GLMsingle algorithm (Allen et al., 2022; Kay et al., 2013; Rokem and Kay, 2020) was used to derive nuisance regressors and to choose the optimum ridge regularization shrinkage fraction for each voxel. The extracted betas for each voxel represent estimates of the trial-wise BOLD response amplitudes to each stimulus trial, and these are relative to the BOLD signal observed during the absence of a stimulus (when only the grey screen was shown). Trials showing the same image were averaged to improve signal estimates and reduce the amount of data. All analyses were done in MATLAB 2019a (MathWorks Inc., Natick, USA).

### Color image statistics

The RGB images were converted to LMS cone tristimulus values using the 10 degree Stockman, MacLeod, Johnson cone fundamentals (Stockman et al., 1993) interpolated to 1nm. Chromaticity coordinates in a version of the MacLeod-Boynton chromaticity diagram (MacLeod and Boynton, 1979) based on the cone fundamentals were extracted for each pixel. In this color diagram, the cardinal mechanisms of color vision are represented by the axes L/(L+M) (roughly teal and red colors) and S/(L+M) (roughly chartreuse to violet), which correspond to the two main retinogeniculate color pathways (Mollon and Cavonius, 1987). Saturation was defined as the distance between the values of the pixel in Macleod-Boynton color space and the NSD grey background. To do this the chromaticity coordinates in the MacLeod-Boynton chromaticity diagram were transformed to polar coordinates (Bosten and Lawrance-Owen, 2014). The scaling factor applied to the L/(L+M) axis was 0.045. If the luminance of a pixel value fell below a dark filtering criterion of L+M = 0.0002, the saturation value was set to zero because at low luminance there is a high level of chromatic noise which would be perceptually very dark or black. The saturation values for each pixel were then averaged over the image to find the average saturation of each image. We used the 425 × 425 images for all analyses of image statistics.

### Definition of ROIs

We created Regions of Interest for the medial and lateral ROIs for both hemispheres. Using the map of the number of participants that showed a whole-brain Bonferroni-corrected significant correlation between voxel response and average saturation for each voxel in fsaverage space (Fig. 1B). For both hemispheres we drew large ROIs around each stream (medial and lateral) of voxel responses that correlated significantly (following a whole-brain Bonferroni correction) with average saturation in at least one participant, beginning at the boundary of Kastner-defined hV4 (Fig. 1B). We applied the four ROIs to each participant but only included voxels in an ROI for a particular participant if the voxels responses showed significant positive correlations with average saturation (again, Bonferroni-corrected over the whole brain).

### Creation of montages

We Z-scored voxel responses to all images for each voxel and then averaged the Z-scored voxel responses across voxels in each ROI for each image. Using the average voxel responses for each ROI we created montages of images that evoked the highest and lowest average voxel responses. We plotted four hundred images in each montage out of the 9,209 - 10,000 images each participant saw. A part of each montage is shown in Figure 2B.

### Other image statistics

For pixel counts of food and circular objects, we summed the number of pixels containing food or circular objects for each image. To do this we used the COCO dataset object segmentation data which has 80 object categories. We converted the segmentation data to a binary pixel mask for each image which included the object, then the total number of pixels was summed. The food image categories were banana, apple, sandwich, orange, broccoli, carrot, hot dog, pizza, donut and cake. Circular object categories were sport ball, pizza, donut, clock, tennis racket, frisbee, wine glass, stop sign, and cup. It is possible that there are other food and/or circular objects in the dataset that were not segmented. For the warm-cool image statistic, we used warm-cool color ratings collected by our group for another project (Maule, Racey, Tang, Richter, Bird & Franklin, unpublished), where participants were shown a set of 24 isoluminant and iso-saturated hues and asked to rate how warm-cool they appeared using sliding scale. We used these warm-cool ratings to interpolate the warm-cool value for the hue of each pixel that had a luminance higher than the dark filter criterion described previously. Warm ratings had positive values and cool ratings had negative values. We summed the warm-cool values of all pixels in the image to get an overall warm-cool statistic for each image.

### Relationships between image statistics and voxel responses

To create Figure 3A, we ranked the images for each image statistic and then averaged over the lowest ranking 500 images (images ranked 1 to 500). We also averaged over the Z-scored voxel responses to the same 500 images. We repeated this procedure but selected images ranking between 2 and 501 and the corresponding voxel responses. We continued moving one image up until reaching the highest ranking 500 images. Afterwards, we extrapolated the resulting “moving-average” curves to the highest and lowest image statistic values seen by any of the 8 participants. We then averaged across the eight subjects at interpolated points along the image statistic. The interpolation was necessary because each subject saw different images (other than the roughly 10% common images). In SI Figure 3, plots for individual participants are shown.

### Multiple linear regression

We applied a rank inverse normal transform (Blom constant) to all image statistics before conducting the multiple regression. Responses for each individual voxel were Z-scored across images and then average voxel responses for each image were calculated for each of the four ROIs.

### ANOVA with saturation and food

The median image average saturation was used to split the saturation variable into high and low saturation individually for each participant. We categorized images that contained food based on the COCO categories and all other images were categorized as non-food images. From these we created four groups of images: high food/low saturation, low food/low saturation, high food/high saturation, and low food/ high saturation. On average, the food group of images was more saturated (M = 0.7462, SD = 0.0437) than the non-food group (M = 0.4793, SD = 0.0382). A Welch’s t-test showed there was a significant difference between the groups for all eight participants (24.74 ≤ t(1541.5) ≤ 26.40, 2.78 × 10^−113^ ≥ *p* ≥ 1.16 × 10^−127^ across 8 participants).

## Acknowledgements

Collection and pre-processing of MRI data was supported by NSF IIS-1822683 (K.K.), NSF IIS-1822929 (T.N.), NIH S10 RR026783, the W.M. Keck Foundation, the analyses described here by European Research Council grants COLOURMIND 772193 (AF) and COLOURCODE 949242 (JB)

