## Supplementary Material for "Color-biased regions in the ventral visual pathway are food-selective"

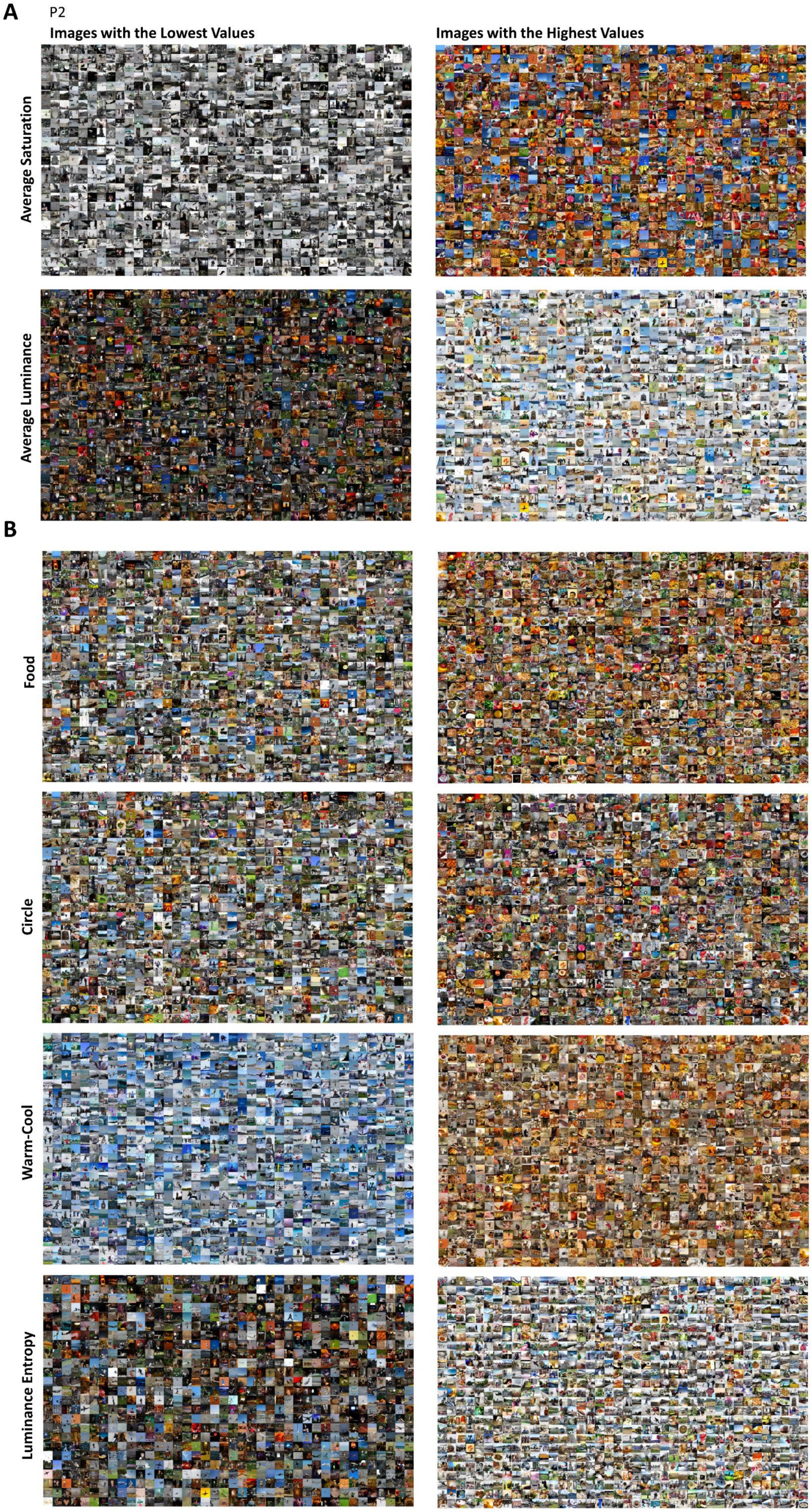

A
P2
Images with the Lowest Values
Images with the Highest Values
Average Saturation
Average Luminance
B
Food
Circle
Warm-Cool
Luminance Entropy




2

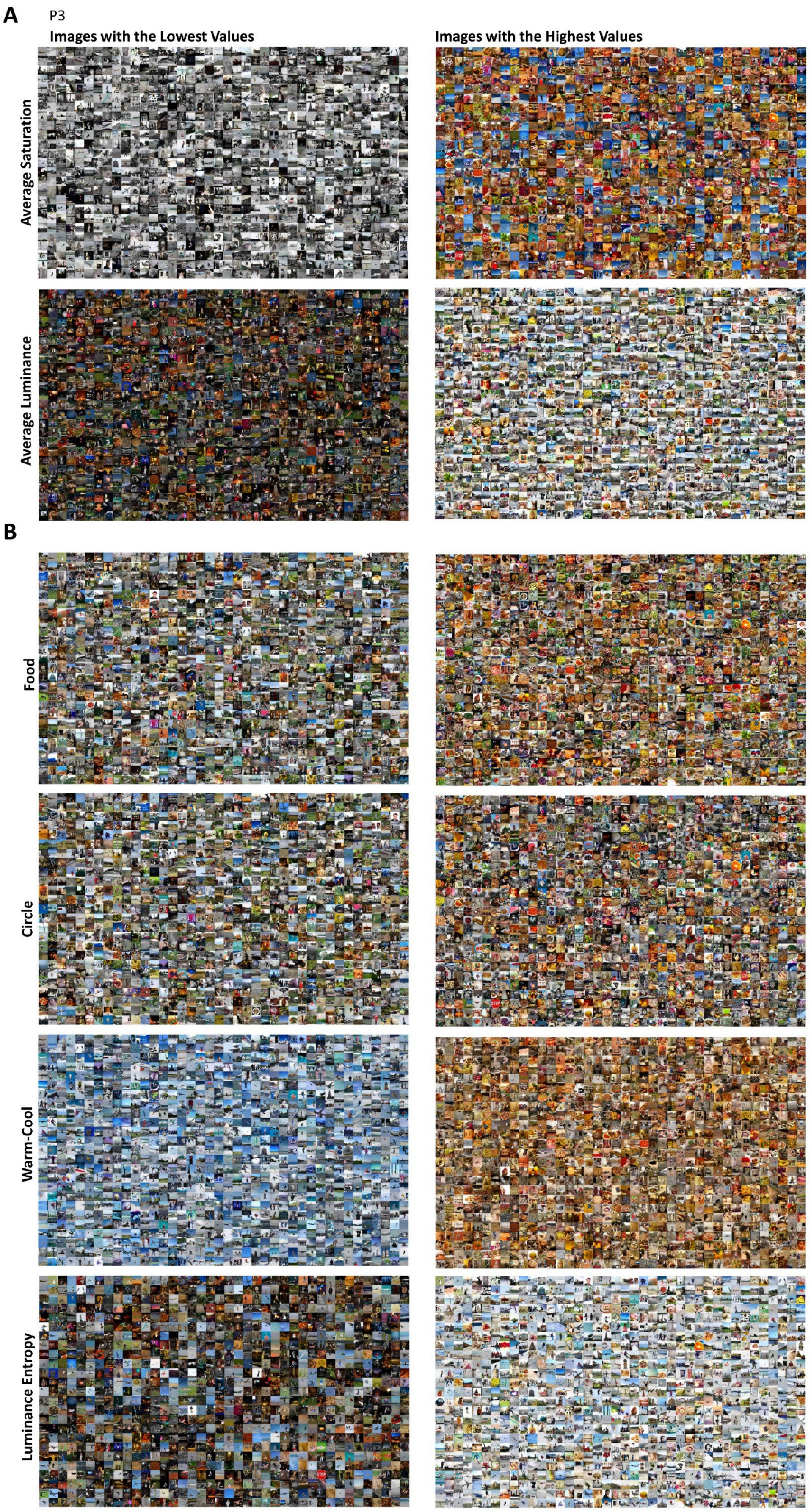

A
P3
Images with the Lowest Values
Images with the Highest Values
Average Saturation
Average Luminance
B
Food
Circle
Warm-Cool
Luminance Entropy




3

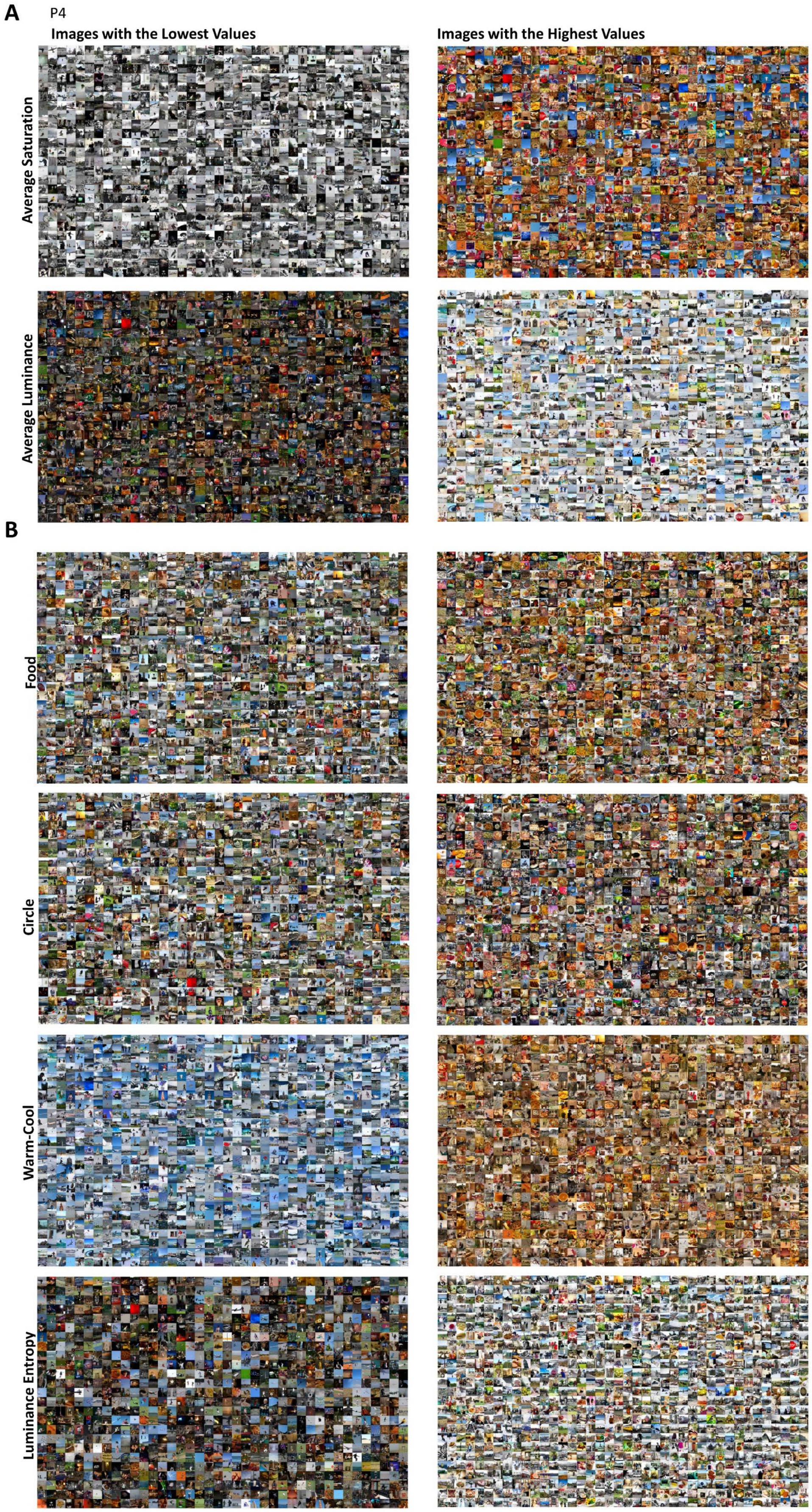
A
P4
Images with the Lowest Values
Images with the Highest Values
Average Saturation
Average Luminance
B
Food
Circle
Warm-Cool
Luminance Entropy



4

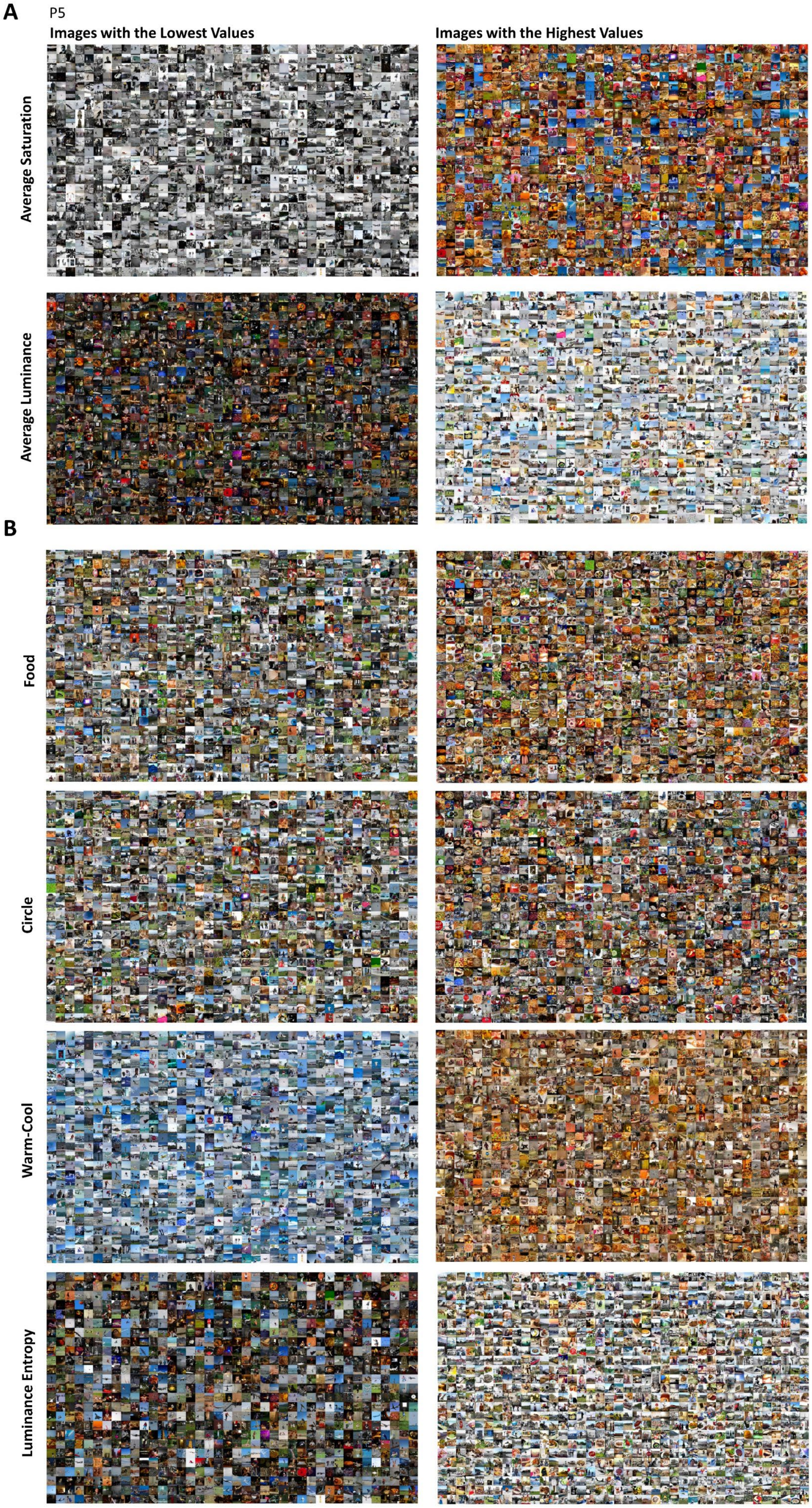
A
P5
Images with the Lowest Values
Images with the Highest Values
Average Saturation
Average Luminance
B
Food
Circle
Warm-Cool
Luminance Entropy



5

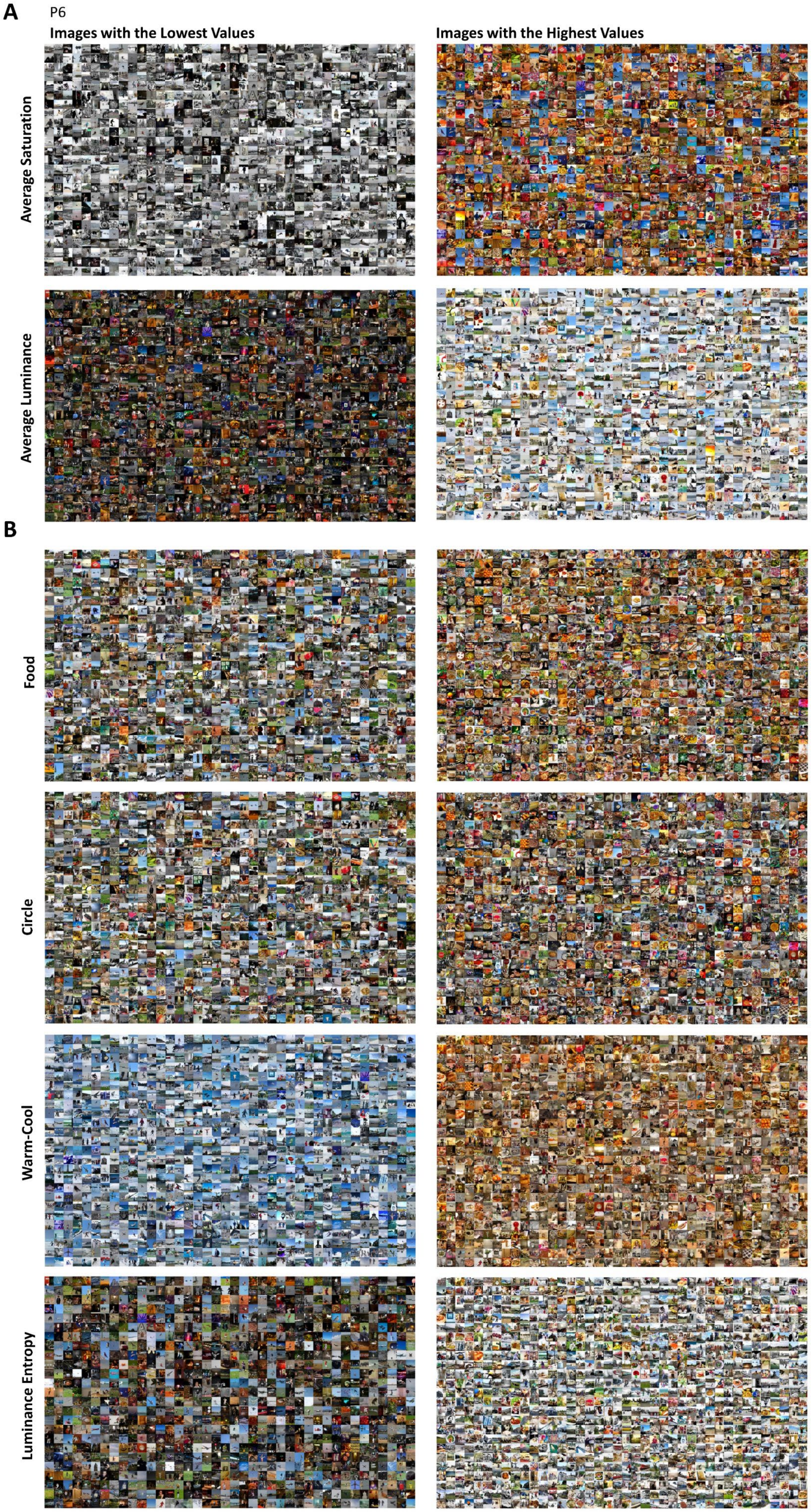
A
P6
Images with the Lowest Values
Images with the Highest Values
Average Saturation
Average Luminance
B
Food
Circle
Warm-Cool
Luminance Entropy


6

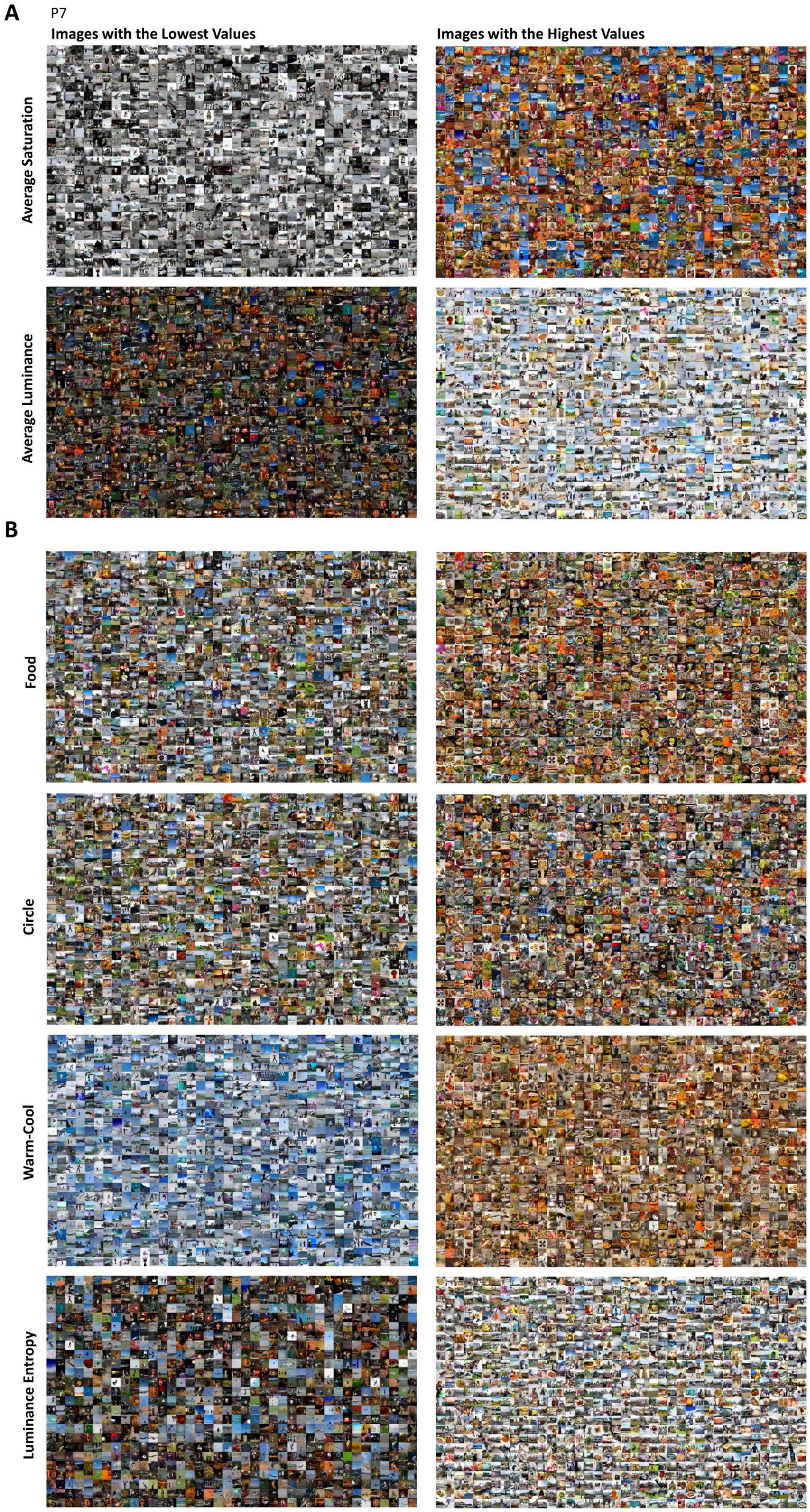
A
P7
Images with the Lowest Values
Images with the Highest Values
Average Saturation
Average Luminance
B
Food
Circle
Warm-Cool
Luminance Entropy

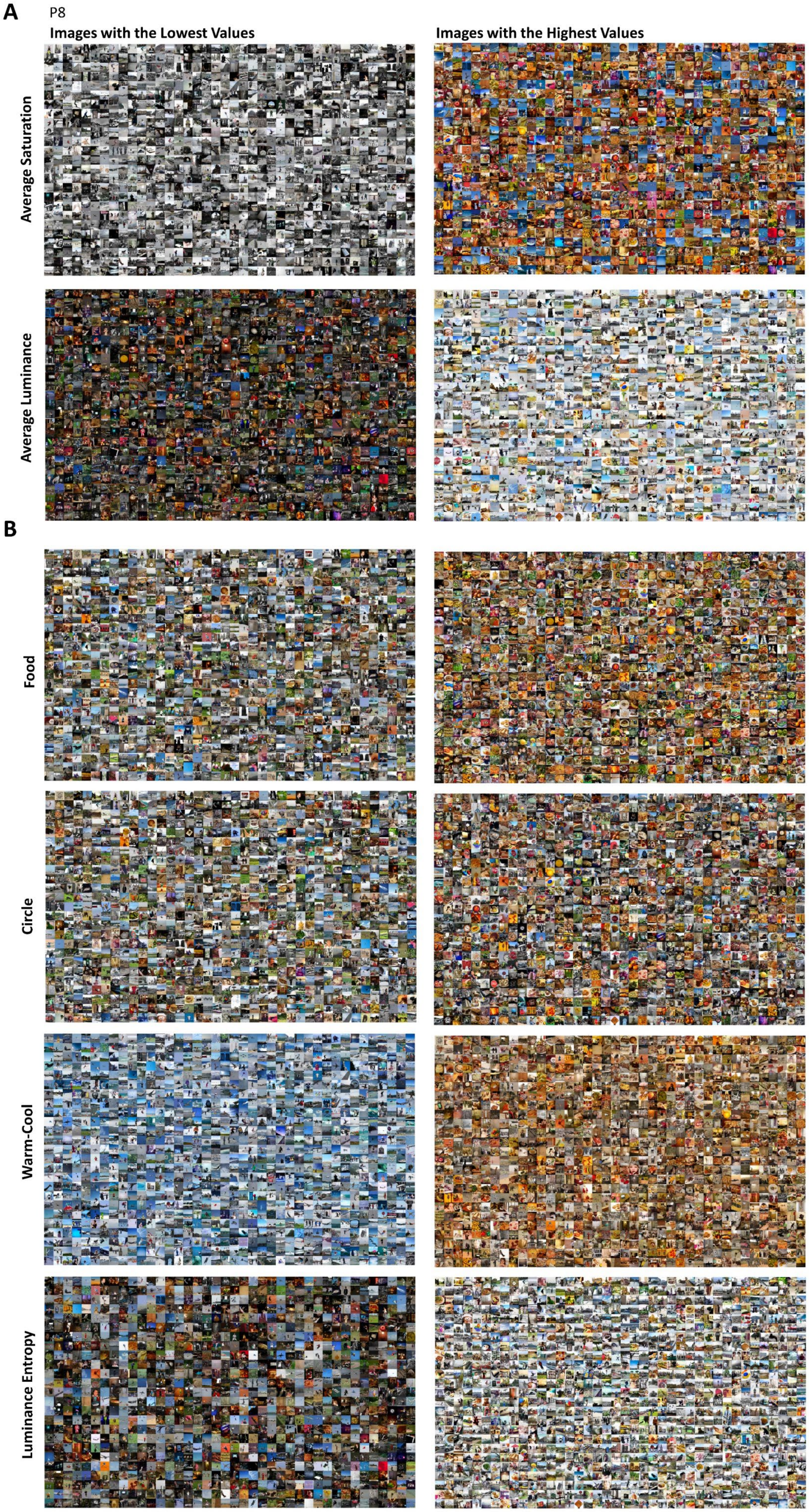
A
P8
Images with the Lowest Values
Images with the Highest Values
Average Saturation
Average Luminance
B
Food
Circle
Warm-Cool
Luminance Entropy



8

**Figure 1: Montages of lowest and highest images for each image statistic**

Montages of the thousand images with lowest (left) and highest (right) image statistics for participants 2 to 8. (A) Montages of the image statistics included in the correlation analysis: average saturation (based on the NSD grey background), and average luminance which was a covariate. (B) For other analyses four further image statistics were added: food pixel count, circle pixel count, warm-cool ratings and luminance entropy.

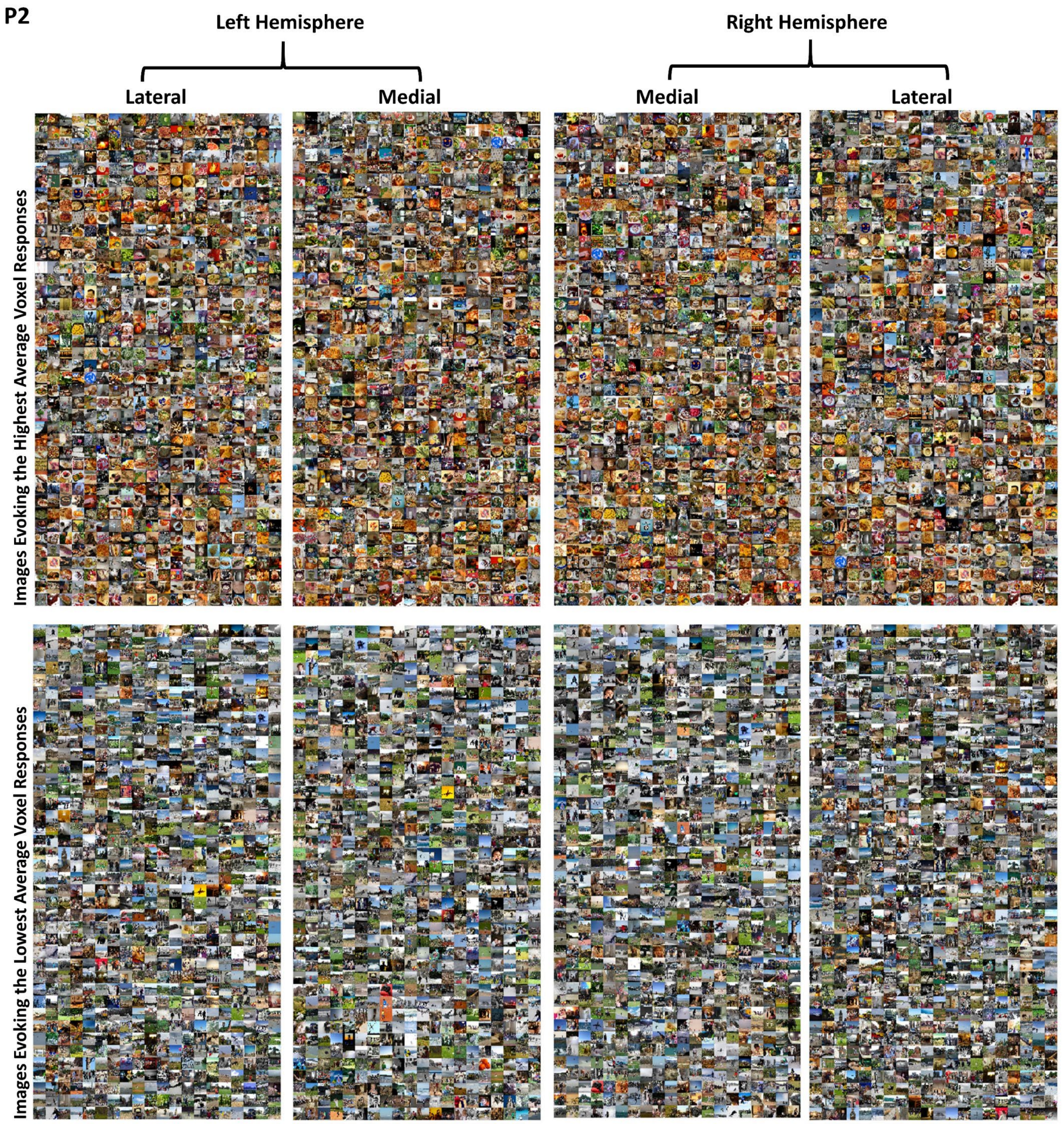
P2
Left Hemisphere
Right Hemisphere
Lateral
Medial
Medial
Lateral
Images Evoking the Highest Average Voxel Responses
Images Evoking the Lowest Average Voxel Responses

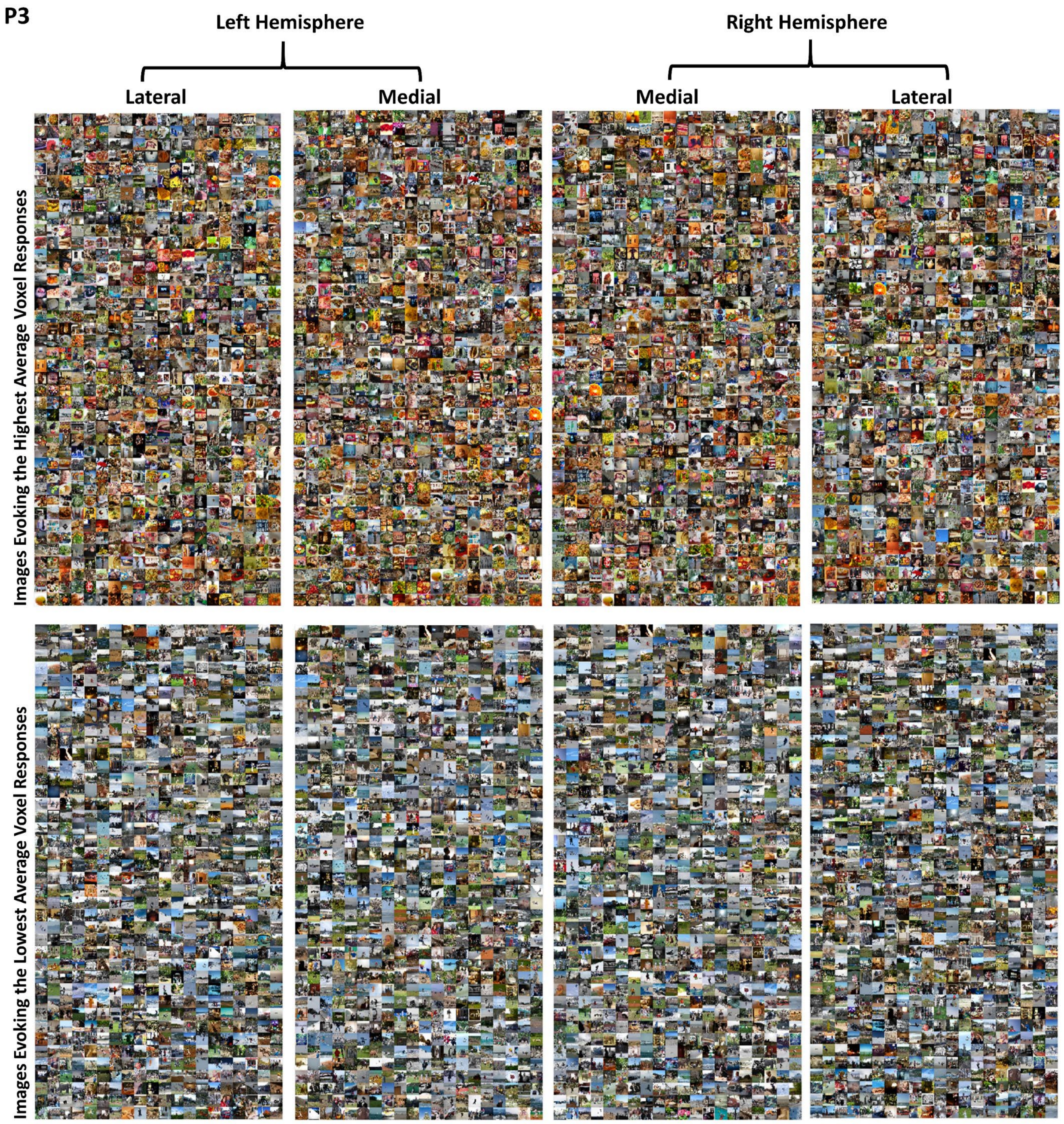
P3
Left Hemisphere
Right Hemisphere
Lateral
Medial
Medial
Lateral
Images Evoking the Highest Average Voxel Responses
Images Evoking the Lowest Average Voxel Responses

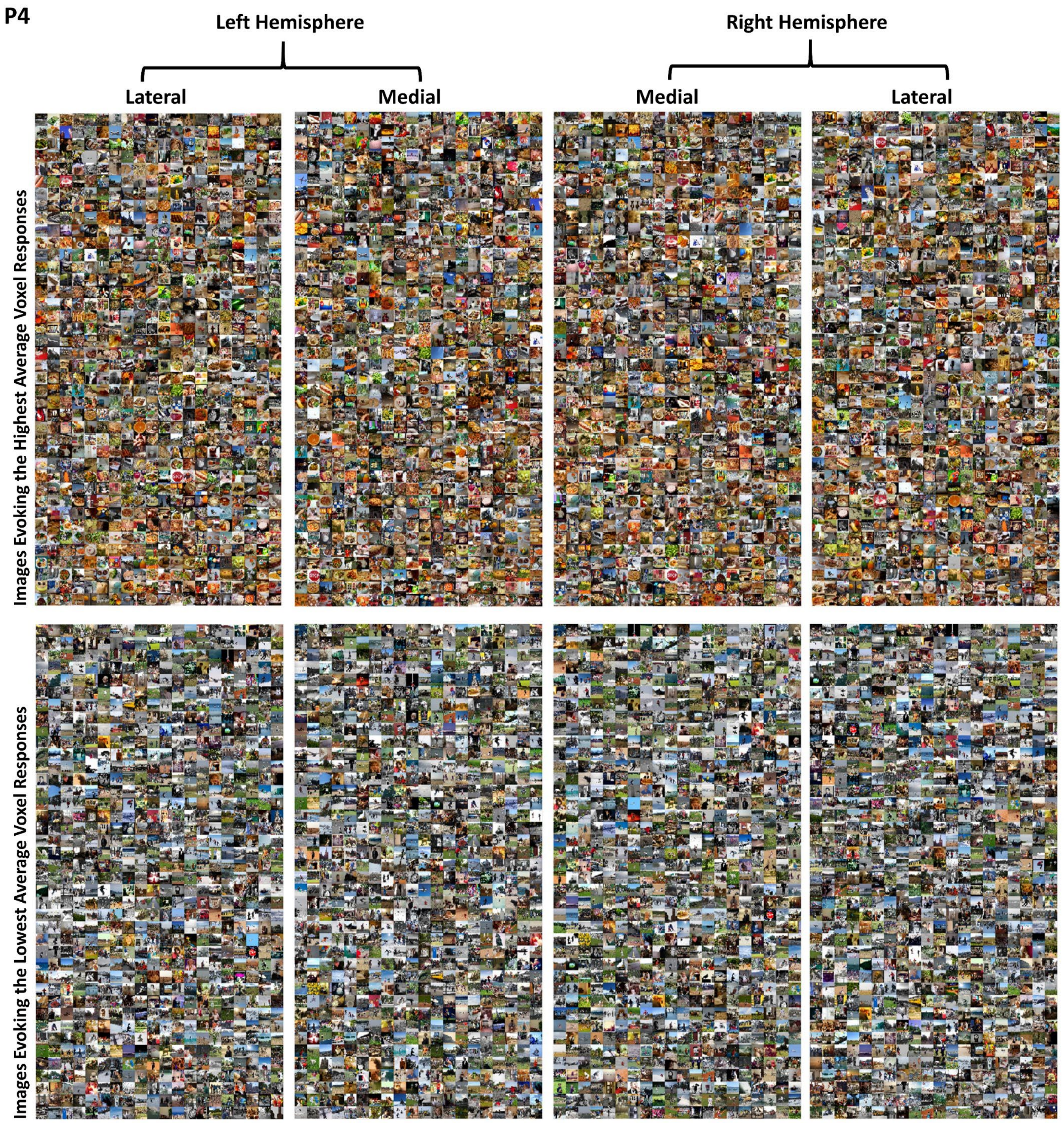
P4
Left Hemisphere
Right Hemisphere
Lateral
Medial
Medial
Lateral
Images Evoking the Highest Average Voxel Responses
Images Evoking the Lowest Average Voxel Responses

17

P5

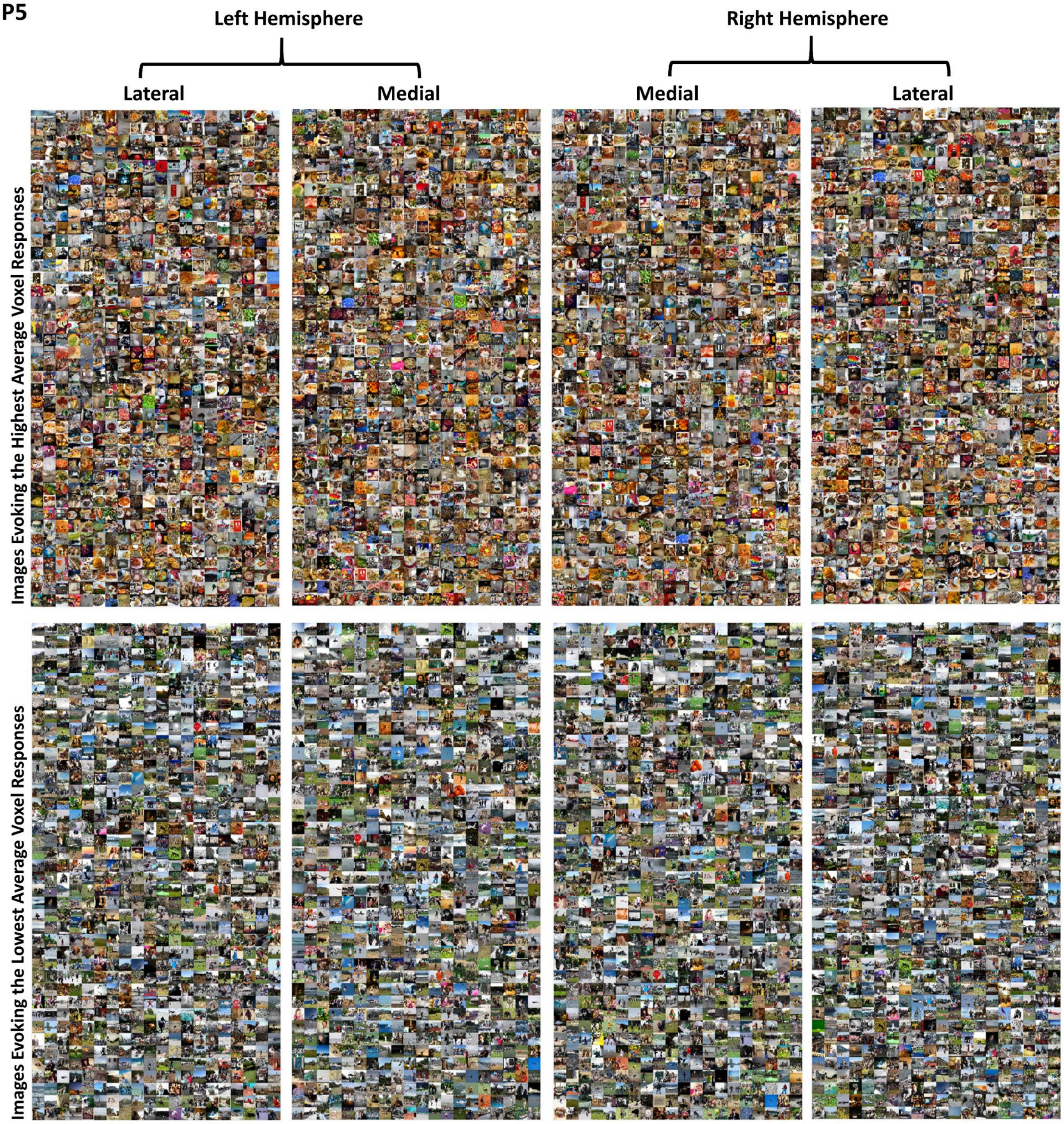
Left Hemisphere
Right Hemisphere
Lateral
Medial
Medial
Lateral
Images Evoking the Highest Average Voxel Responses
Images Evoking the Lowest Average Voxel Responses

18

P6

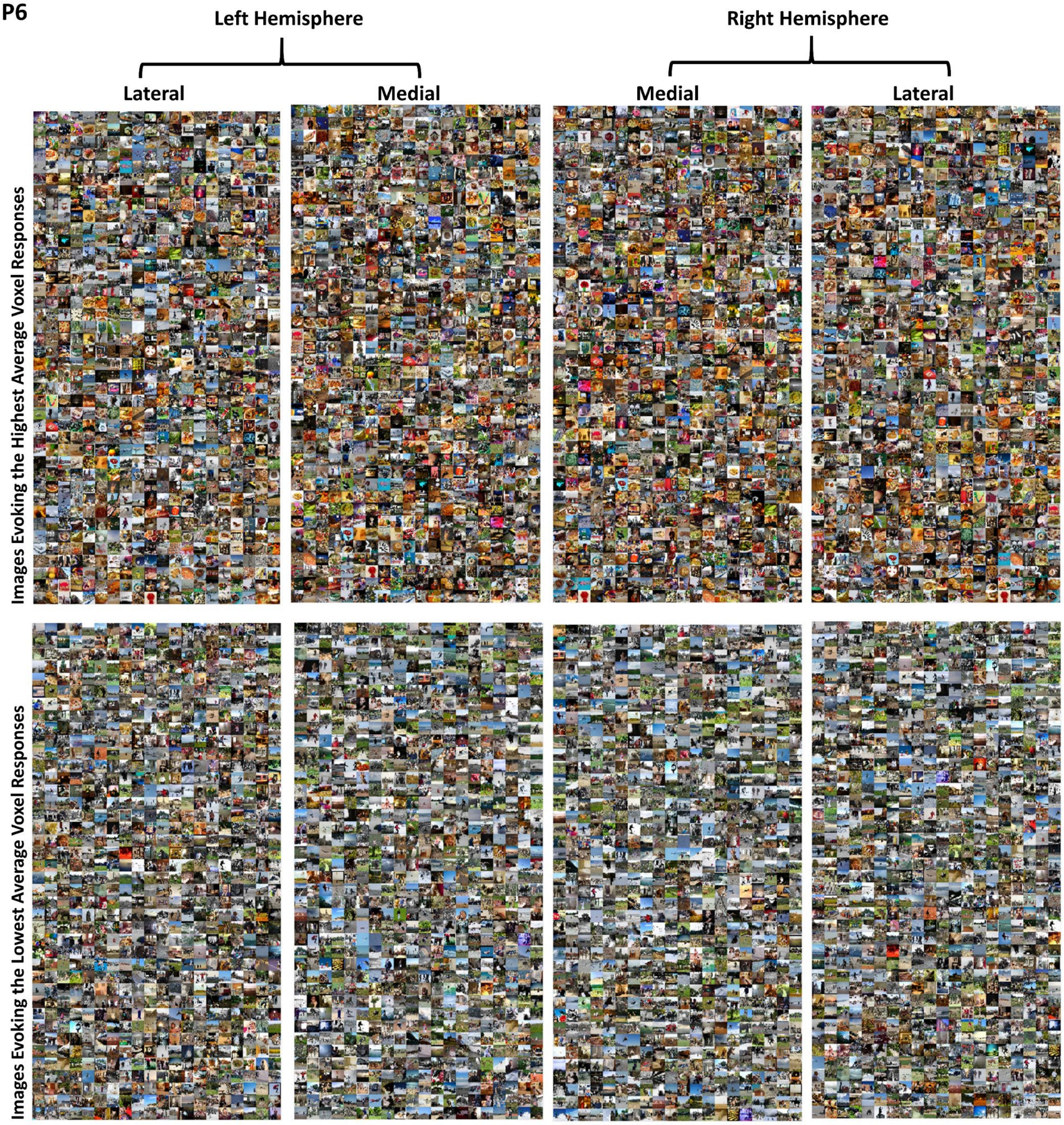
Left Hemisphere
Right Hemisphere
Lateral
Medial
Medial
Lateral
Images Evoking the Highest Average Voxel Responses
Images Evoking the Lowest Average Voxel Responses

19

**P7**

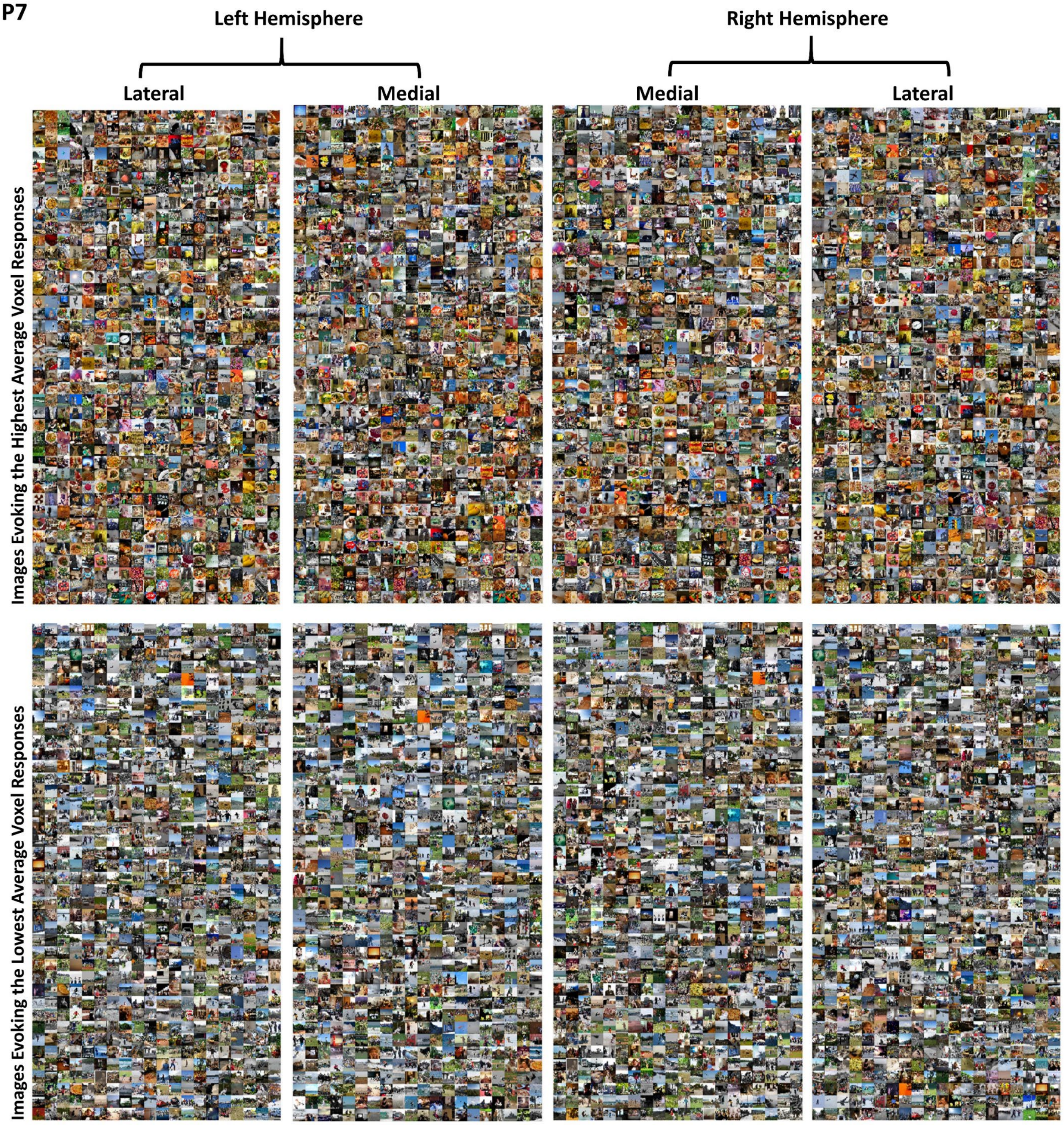
Left Hemisphere
Right Hemisphere
Lateral
Medial
Medial
Lateral
Images Evoking the Highest Average Voxel Responses
Images Evoking the Lowest Average Voxel Responses

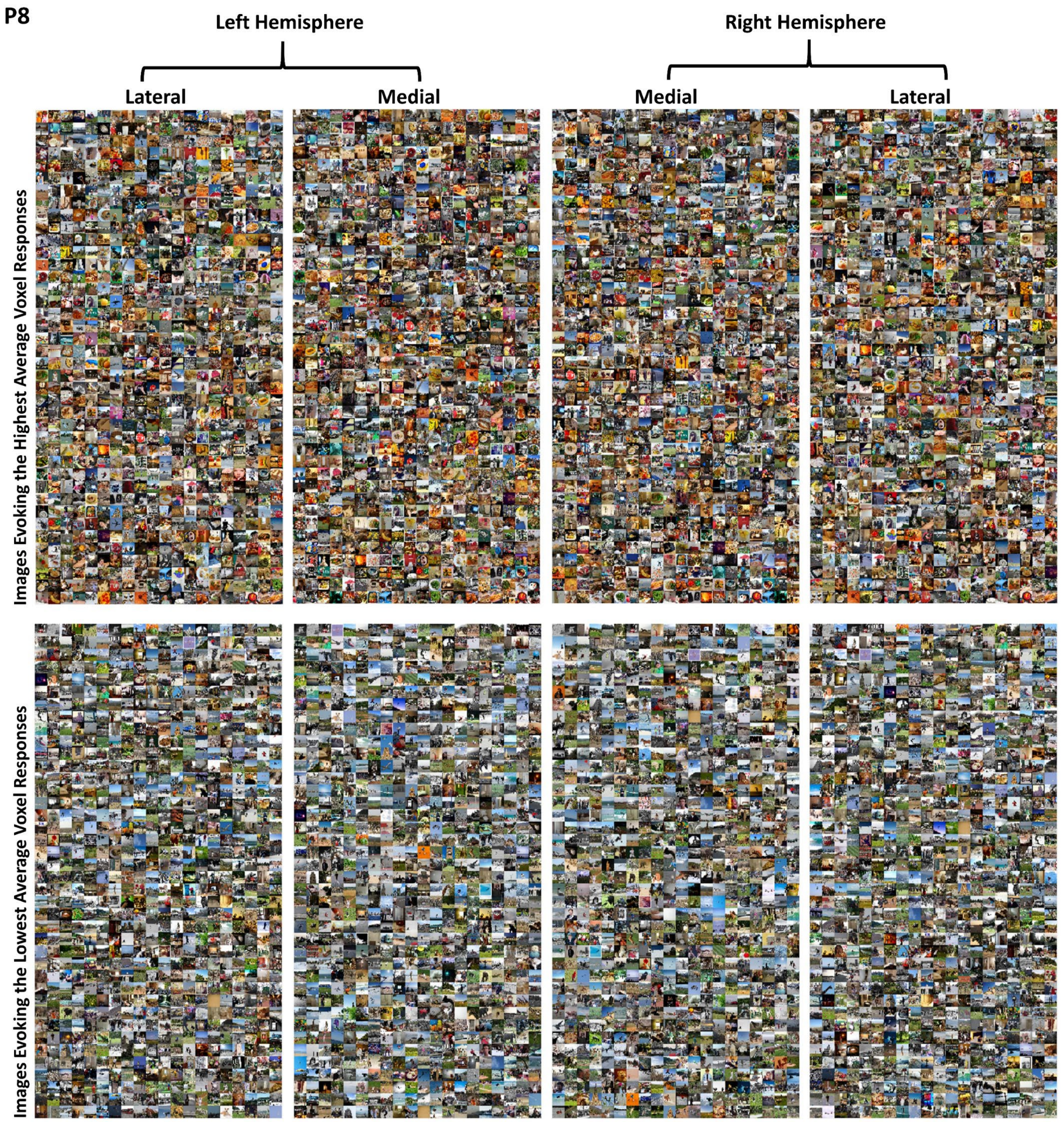


**Figure 2: Montages of image evoking the highest and lowest average voxel responses in the four ROIs.**

Montages show the images for the lateral and medial areas for both hemispheres that provoke the highest and lowest averaged Z-scored voxel responses for participants two to eight. The 800 images evoking the highest and lowest voxel responses were selected for each montage (from left to right: left hemisphere-lateral ROI, left hemisphere-medial ROI, right hemisphere-medial ROI, right hemisphere-lateral ROI. The upper row shows images evoking the highest responses, and the lower row shows images evoking the lowest responses.

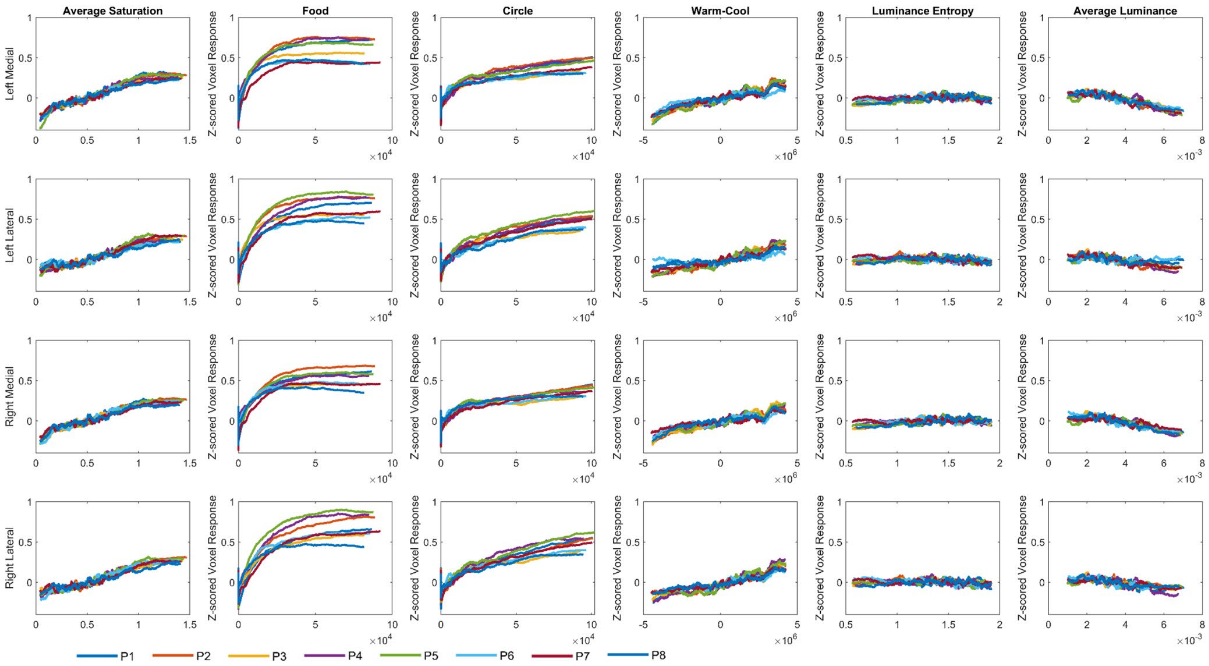


**Figure 3: Average ROI responses for each of the image statistics**

Each row shows the interpolated average Z-scored voxel responses across each ROI, with data from each of the eight participants plotted in a different color. The x-axis shows an image statistic: average saturation, number of pixels contained in images of food, number of pixels contain in images of circular objects, summed warm-cool ratings based on individual pixels, luminance (L+M) entropy (Mather, 2020), and average luminance. The y-axis shows the averaged Z-scored voxel responses. The images were ranked from lowest to highest for each image statistic and the image statistic was averaged over sequential sets of 500. Then the average of 500 Z-scored voxel responses were sequentially plotted for the corresponding sets of 500 images, e.g., 1-500, 2-501, 3-502, creating a “moving average” voxel response for each ROI as each image statistic increased.

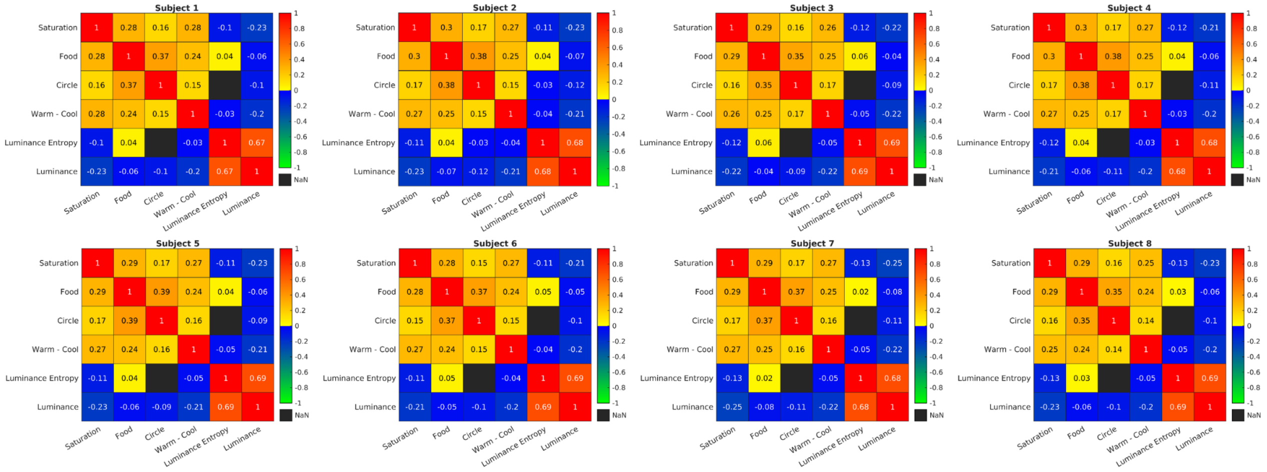


**Figure 4: Inter-correlations between image statistics for each participant**

These correlations show the relationships between our six image statistics for each participant. All pairs of image statistics were significantly correlated apart for luminance (L+M) entropy and circle pixel count which were significantly correlated for participant 2 but not for the other 7 participants.

| | P | Sum of Squared | | | | | df | | | | | Mean Squared | | | | F | | | Prob>F | | |
|---|---|---|---|---|---|---|---|---|---|---|---|---|---|---|---|---|---|---|---|---|---|
| | | Food | Saturation | Food*Saturation | Error | Total | Food | Saturation | Food*Saturation | Error | Total | Food | Saturation | Food*Saturation | Error | Food | Saturation | Food*Saturation | Food | Saturation | Food*Saturation |
| Left | 1 | 205.4707 | 28.42099 | 4.7989502 | 2695 | 3132.5 | 1 | 1 | 1 | 9996 | 9999 | 205.5 | 28.42099 | 4.7989502 | 0.27 | 762.03 | 105.40471 | 17.797831 | 0 | 1.32E-24 | 2.48E-05 |
| Medial | 2 | 208.5696 | 16.178802 | 3.6601868 | 2366 | 2819.9 | 1 | 1 | 1 | 9996 | 9999 | 208.6 | 16.178802 | 3.6601868 | 0.237 | 881.35 | 68.366348 | 15.466758 | 0 | 1.53E-16 | 8.45E-05 |
| | 3 | 119.04 | 15.226837 | 2.2487793 | 1852 | 2146.5 | 1 | 1 | 1 | 9407 | 9410 | 119 | 15.226837 | 2.2487793 | 0.197 | 604.64 | 77.341263 | 11.422164 | 0 | 1.69E-18 | 0.000728681 |
| | 4 | 197.7175 | 17.905029 | 3.6851807 | 2853 | 3273.1 | 1 | 1 | 1 | 9205 | 9208 | 197.7 | 17.905029 | 3.6851807 | 0.31 | 637.87 | 57.76424 | 11.888931 | 0 | 3.24E-14 | 0.000567212 |
| | 5 | 220.9007 | 15.976898 | 0.88864136 | 2664 | 3115.6 | 1 | 1 | 1 | 9996 | 9999 | 220.9 | 15.976898 | 0.88864136 | 0.267 | 828.78 | 59.9422 | 3.3340087 | 0 | 1.07E-14 | 0.067891106 |
| | 6 | 114.8306 | 7.4012146 | 0.046585083 | 2491 | 2706.4 | 1 | 1 | 1 | 9407 | 9410 | 114.8 | 7.4012146 | 0.046585083 | 0.265 | 433.62 | 27.948185 | 0.17591283 | 0 | 1.27E-07 | 0.67491907 |
| | 7 | 86.20985 | 9.6344604 | 0.19499207 | 3000 | 3180.9 | 1 | 1 | 1 | 9996 | 9999 | 86.21 | 9.6344604 | 0.19499207 | 0.3 | 287.22 | 32.098469 | 0.64964169 | 0 | 1.51E-08 | 0.42025998 |
| | 8 | 81.93811 | 12.69046 | 1.277832 | 2255 | 2458.8 | 1 | 1 | 1 | 9205 | 9208 | 81.94 | 12.69046 | 1.277832 | 0.245 | 334.48 | 51.804501 | 5.2163157 | 0 | 6.61E-13 | 0.022398578 |
| Left | 1 | 163.7947 | 11.81192 | 4.2930603 | 2736 | 3046.6 | 1 | 1 | 1 | 9996 | 9999 | 163.8 | 11.81192 | 4.2930603 | 0.274 | 598.49 | 43.15966 | 15.686443 | 0 | 5.30E-11 | 7.53E-05 |
| Lateral | 2 | 236.9721 | 8.7453003 | 3.2997131 | 2815 | 3285.1 | 1 | 1 | 1 | 9996 | 9999 | 237 | 8.7453003 | 3.2997131 | 0.282 | 841.43 | 31.052303 | 11.71643 | 0 | 2.58E-08 | 0.000621991 |
| | 3 | 123.6994 | 7.0943604 | 0.86090088 | 2277 | 2541.4 | 1 | 1 | 1 | 9407 | 9410 | 123.7 | 7.0943604 | 0.86090088 | 0.242 | 511.01 | 29.307076 | 3.5564146 | 0 | 6.33E-08 | 0.059346512 |
| | 4 | 170.8998 | 20.601624 | 8.2520752 | 2922 | 3309.2 | 1 | 1 | 1 | 9205 | 9208 | 170.9 | 20.601624 | 8.2520752 | 0.317 | 538.32 | 64.893456 | 25.993372 | 0 | 8.89E-16 | 3.49E-07 |
| | 5 | 300.1193 | 8.2953491 | 0.98516846 | 3150 | 3694 | 1 | 1 | 1 | 9996 | 9999 | 300.1 | 8.2953491 | 0.98516846 | 0.315 | 952.29 | 26.321442 | 3.1259751 | 0 | 2.94E-07 | 0.077084258 |
| | 6 | 103.5693 | 4.0707855 | 1.2955627 | 3744 | 3925.1 | 1 | 1 | 1 | 9407 | 9410 | 103.6 | 4.0707855 | 1.2955627 | 0.398 | 260.21 | 10.227651 | 3.2550385 | 0 | 0.00138809 | 0.071236238 |
| | 7 | 133.2122 | 10.084137 | 0.33346558 | 3362 | 3620.3 | 1 | 1 | 1 | 9996 | 9999 | 133.2 | 10.084137 | 0.33346558 | 0.336 | 396.02 | 29.978504 | 0.99133909 | 0 | 4.47E-08 | 0.31943941 |
| | 8 | 71.86323 | 7.9950409 | 0.86013794 | 2928 | 3093.3 | 1 | 1 | 1 | 9205 | 9208 | 71.86 | 7.9950409 | 0.86013794 | 0.318 | 225.95 | 25.137299 | 2.7043695 | 0 | 5.44E-07 | 0.10010778 |
| Right | 1 | 134.887 | 21.381866 | 3.7539368 | 2383 | 2678.7 | 1 | 1 | 1 | 9996 | 9999 | 134.9 | 21.381866 | 3.7539368 | 0.238 | 565.9 | 89.704056 | 15.749016 | 0 | 3.39E-21 | 7.28E-05 |
| Medial | 2 | 158.1331 | 17.460388 | 3.5646362 | 1786 | 2153.3 | 1 | 1 | 1 | 9996 | 9999 | 158.1 | 17.460388 | 3.5646362 | 0.179 | 885.01 | 97.718994 | 19.949881 | 0 | 6.14E-23 | 8.04E-06 |
| | 3 | 75.10976 | 14.75708 | 0.71833801 | 2583 | 2793.3 | 1 | 1 | 1 | 9407 | 9410 | 75.11 | 14.75708 | 0.71833801 | 0.275 | 273.51 | 53.738335 | 2.6158488 | 0 | 2.48E-13 | 0.10583443 |
| | 4 | 113.9371 | 9.7249451 | 1.584671 | 2112 | 2351 | 1 | 1 | 1 | 9205 | 9208 | 113.9 | 9.7249451 | 1.584671 | 0.229 | 496.61 | 42.387627 | 6.9070258 | 0 | 7.88E-11 | 0.00860002 |
| | 5 | 144.1196 | 11.05542 | 0.98306274 | 2511 | 2808.3 | 1 | 1 | 1 | 9996 | 9999 | 144.1 | 11.05542 | 0.98306274 | 0.251 | 573.69 | 44.007904 | 3.9132419 | 0 | 3.44E-11 | 0.047934514 |
| | 6 | 107.1402 | 13.971252 | 0.008407593 | 2440 | 2671.7 | 1 | 1 | 1 | 9407 | 9410 | 107.1 | 13.971252 | 0.008407593 | 0.259 | 413.04 | 53.861473 | 0.032412652 | 0 | 2.33E-13 | 0.85712886 |
| | 7 | 85.73183 | 9.6898956 | 0.56030273 | 2859 | 3038 | 1 | 1 | 1 | 9996 | 9999 | 85.73 | 9.6898956 | 0.56030273 | 0.286 | 299.72 | 33.876591 | 1.9588597 | 0 | 6.05E-09 | 0.16166636 |
| | 8 | 61.23292 | 7.9783173 | 0.38876343 | 2803 | 2950.4 | 1 | 1 | 1 | 9205 | 9208 | 61.23 | 7.9783173 | 0.38876343 | 0.305 | 201.06 | 26.197567 | 1.2765418 | 4.20E-45 | 3.14E-07 | 0.25857246 |
| Right | 1 | 121.9491 | 12.04422 | 2.8839417 | 3302 | 3544.7 | 1 | 1 | 1 | 9996 | 9999 | 121.9 | 12.04422 | 2.8839417 | 0.33 | 369.16 | 36.459919 | 8.7301865 | 0 | 1.61E-09 | 0.003137143 |
| Lateral | 2 | 191.5377 | 12.939056 | 4.340271 | 2847 | 3257.5 | 1 | 1 | 1 | 9996 | 9999 | 191.5 | 12.939056 | 4.340271 | 0.285 | 672.48 | 45.42873 | 15.238592 | 0 | 1.67E-11 | 9.54E-05 |
| | 3 | 82.29128 | 11.567902 | 2.2331848 | 3186 | 3393.8 | 1 | 1 | 1 | 9407 | 9410 | 82.29 | 11.567902 | 2.2331848 | 0.339 | 242.98 | 34.156624 | 6.5939403 | 0 | 5.25E-09 | 0.010247947 |
| | 4 | 209.2648 | 17.756348 | 5.9180603 | 3838 | 4280.5 | 1 | 1 | 1 | 9205 | 9208 | 209.3 | 17.756348 | 5.9180603 | 0.417 | 501.83 | 42.581242 | 14.192016 | 0 | 7.14E-11 | 0.000166099 |
| | 5 | 326.1877 | 10.0672 | 1.3817139 | 2974 | 3571.2 | 1 | 1 | 1 | 9996 | 9999 | 326.2 | 10.0672 | 1.3817139 | 0.297 | 1096.5 | 33.842567 | 4.6448607 | 0 | 6.16E-09 | 0.031170467 |
| | 6 | 151.3702 | 12.077698 | 0.49386597 | 3165 | 3455.1 | 1 | 1 | 1 | 9407 | 9410 | 151.4 | 12.077698 | 0.49386597 | 0.336 | 449.84 | 35.892609 | 1.4676752 | 0 | 2.16E-09 | 0.22574328 |
| | 7 | 119.3219 | 9.1988678 | 2.1040955 | 3089 | 3320.2 | 1 | 1 | 1 | 9996 | 9999 | 119.3 | 9.1988678 | 2.1040955 | 0.309 | 386.16 | 29.770105 | 6.8094401 | 0 | 4.98E-08 | 0.009081231 |
| | 8 | 76.39191 | 7.0662384 | 0.56330872 | 3073 | 3242.6 | 1 | 1 | 1 | 9205 | 9208 | 76.39 | 7.0662384 | 0.56330872 | 0.334 | 228.84 | 21.167784 | 1.6874604 | 0 | 4.26E-06 | 0.19396862 |

**Table 1: Results of a two-way ANOVA with food and saturation as factors for all ROIs and all participants.**

We conducted two-way ANOVAs with food and saturation as factors on average Z-scored voxel responses for all four ROIs. Here we show the results of each individual ANOVA for the individual ROIs and individual participants.

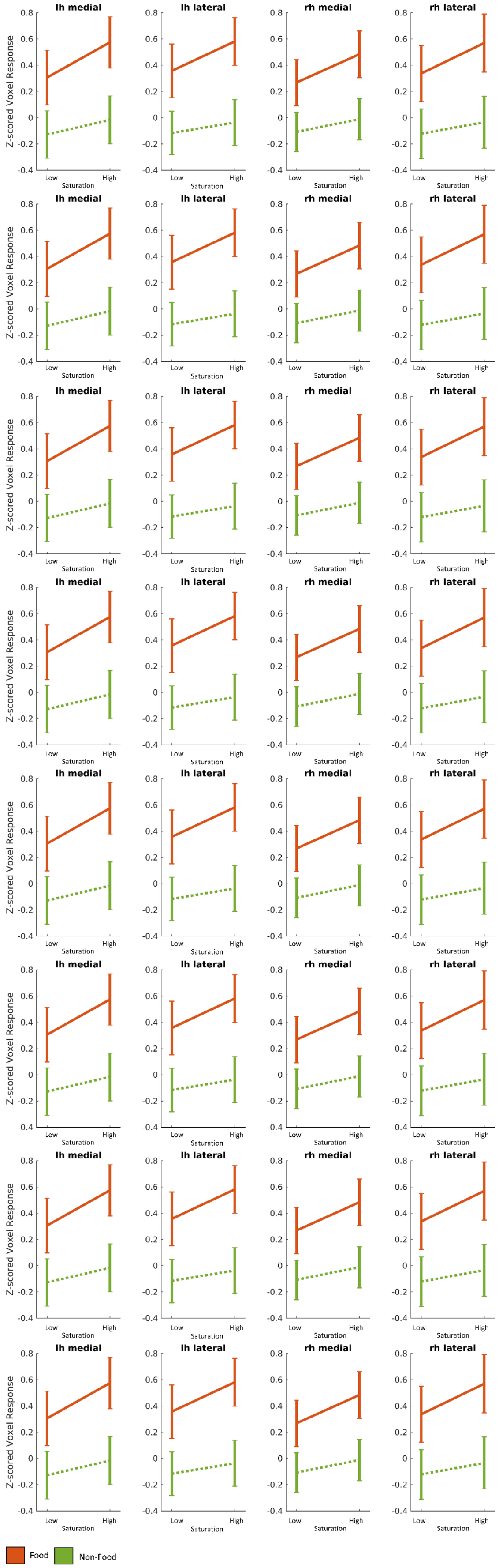
lh medial
lh lateral
rh medial
rh lateral
Z-scored Voxel Response
0.8
0.6
0.4
0.2
0
-0.2
-0.4
Low
High
Saturation
Food
Non-Food

**Figure 5: Two-way ANOVA with food and saturation as factors**

A two-way ANOVA with food and saturation as factors was conducted on average Z-scored voxel response for all four ROIs for each participant. On the x-axis, the low and high saturation image groups are displayed. The y-axis displays the z-scored average voxel responses. The orange line represents results for images that contained food and the green line represents results for images that do not contain food based on the COCO object categories. From top to bottom, participant 1 to 8. Error bars are the 95% confidence intervals.

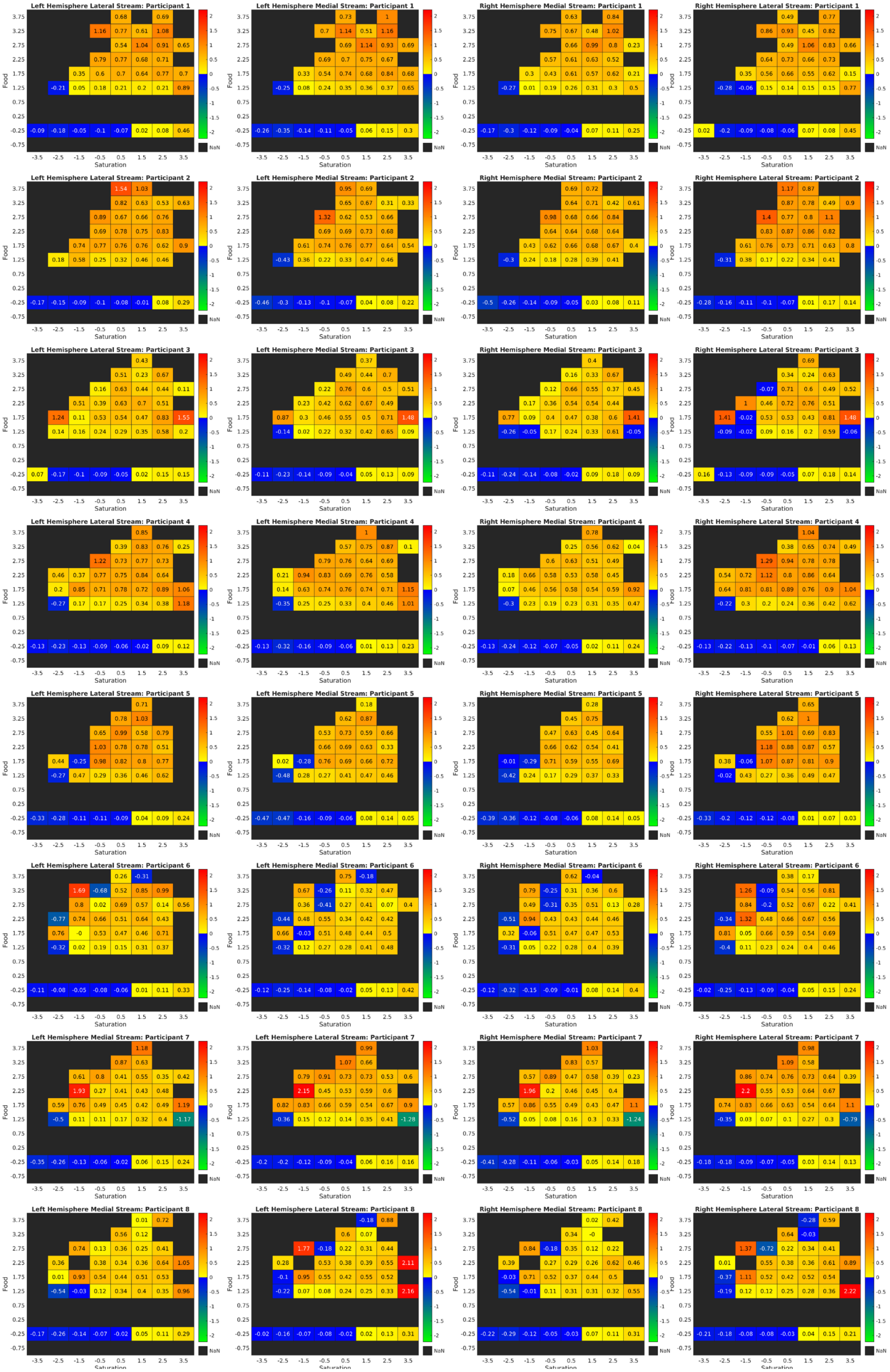

Left Hemisphere Lateral Stream: Participant 1
Left Hemisphere Medial Stream: Participant 1
Right Hemisphere Medial Stream: Participant 1
Right Hemisphere Lateral Stream: Participant 1
Left Hemisphere Lateral Stream: Participant 2
Left Hemisphere Medial Stream: Participant 2
Right Hemisphere Medial Stream: Participant 2
Right Hemisphere Lateral Stream: Participant 2
Left Hemisphere Lateral Stream: Participant 3
Left Hemisphere Medial Stream: Participant 3
Right Hemisphere Medial Stream: Participant 3
Right Hemisphere Lateral Stream: Participant 3
Left Hemisphere Lateral Stream: Participant 4
Left Hemisphere Medial Stream: Participant 4
Right Hemisphere Medial Stream: Participant 4
Right Hemisphere Lateral Stream: Participant 4
Left Hemisphere Lateral Stream: Participant 5
Left Hemisphere Medial Stream: Participant 5
Right Hemisphere Medial Stream: Participant 5
Right Hemisphere Lateral Stream: Participant 5
Left Hemisphere Lateral Stream: Participant 6
Left Hemisphere Medial Stream: Participant 6
Right Hemisphere Medial Stream: Participant 6
Right Hemisphere Lateral Stream: Participant 6
Left Hemisphere Medial Stream: Participant 7
Left Hemisphere Lateral Stream: Participant 7
Right Hemisphere Medial Stream: Participant 7
Right Hemisphere Lateral Stream: Participant 7
Left Hemisphere Medial Stream: Participant 8
Left Hemisphere Lateral Stream: Participant 8
Right Hemisphere Medial Stream: Participant 8
Right Hemisphere Lateral Stream: Participant 8
Food
Saturation
NaN

**Figure 6: Voxel responses for combinations of food and saturation image statistics**

In each cell is the average voxel response for a combination of food (y) and saturation (x) image statistics, with separate heat maps for each of the four ROIs and the 8 participants. Voxel responses were Z-scored, and the image statistics were normalized using a rank inverse normal transform (Blom constant). The x- and y – axes show the bin centers.

**Table 2.** Multiple Linear Regression Beta coefficients

| P1 | B0 | Average Saturation | Food | Circles | Warm-Cool | Luminance Entropy | Average Luminance |
|---|---|---|---|---|---|---|---|
| Medial Left | $-9.9626 \times 10^{-9}$ | 0.0667 | 0.2507 | 0.0500 | 0.0280 | 0.0464 | -0.0547 |
| Medial Right | $4.6606 \times 10^{-9}$ | 0.0324 | 0.2486 | 0.0454 | 0.0148 | -0.0263 | 0.0224 |
| Lateral Left | $-3.7794 \times 10^{-9}$ | 0.0573 | 0.1973 | 0.0535 | 0.0268 | 0.0452 | -0.0484 |
| Lateral Right | $6.4530 \times 10^{-9}$ | 0.0365 | 0.1970 | 0.0496 | 0.0476 | -0.0013 | -0.0037 |
| **P2** | **B0** | **Average Saturation** | **Food** | **Circles** | **Warm-Cool** | **Luminance Entropy** | **Average Luminance** |
| Medial Left | $1.7282 \times 10^{-9}$ | 0.0462 | 0.2580 | 0.0545 | 0.0430 | 0.0451 | -0.0481 |
| Medial Right | $1.0421 \times 10^{-9}$ | 0.0186 | 0.3055 | 0.0248 | 0.0462 | -0.0042 | -0.0133 |
| Lateral Left | $4.4754 \times 10^{-9}$ | 0.0511 | 0.2361 | 0.0335 | 0.0406 | 0.0465 | -0.0447 |
| Lateral Right | $2.2469 \times 10^{-9}$ | 0.0360 | 0.3018 | 0.0028 | 0.0432 | -0.0260 | 0.0222 |
| **P3** | **B0** | **Average Saturation** | **Food** | **Circles** | **Warm-Cool** | **Luminance Entropy** | **Average Luminance** |
| Medial Left | $7.2810 \times 10^{-9}$ | 0.0529 | 0.2021 | 0.0343 | 0.0518 | 0.0480 | -0.0515 |
| Medial Right | $2.7417 \times 10^{-10}$ | 0.0349 | 0.2150 | 0.0222 | 0.0473 | 0.0177 | -0.0263 |
| Lateral Left | $2.1672 \times 10^{-8}$ | 0.0615 | 0.1382 | 0.0430 | 0.0630 | 0.0511 | -0.0592 |
| Lateral Right | $1.7536 \times 10^{-9}$ | 0.0390 | 0.1880 | 0.0166 | 0.0707 | -0.0067 | 0.0006 |
| **P4** | **B0** | **Average Saturation** | **Food** | **Circles** | **Warm-Cool** | **Luminance Entropy** | **Average Luminance** |
| Medial Left | $1.5903 \times 10^{-8}$ | 0.0564 | 0.2724 | 0.0322 | 0.0295 | 0.0190 | -0.0304 |
| Medial Right | $-1.3195 \times 10^{-8}$ | 0.0369 | 0.2815 | 0.0165 | 0.0433 | 0.0068 | -0.0150 |
| Lateral Left | $1.7563 \times 10^{-8}$ | 0.0432 | 0.1944 | 0.0343 | 0.0334 | 0.0253 | -0.0292 |
| Lateral Right | $5.4678 \times 10^{-10}$ | 0.0317 | 0.3056 | 0.0114 | 0.0672 | -0.0113 | -0.0019 |
| **P5** | **B0** | **Average Saturation** | **Food** | **Circles** | **Warm-Cool** | **Luminance Entropy** | **Average Luminance** |
| Medial Left | $1.0096 \times 10^{-8}$ | 0.0713 | 0.2321 | 0.0550 | 0.0547 | 0.0499 | -0.0396 |
| Medial Right | $4.8522 \times 10^{-9}$ | 0.0416 | 0.2999 | 0.0587 | 0.0490 | -0.0073 | 0.0112 |
| Lateral Left | $3.5587 \times 10^{-9}$ | 0.0571 | 0.1873 | 0.0548 | 0.0503 | 0.0330 | -0.0184 |
| Lateral Right | $-6.1675 \times 10^{-9}$ | 0.0364 | 0.3381 | 0.0305 | 0.0578 | -0.0299 | 0.0297 |
| **P6** | **B0** | **Average Saturation** | **Food** | **Circles** | **Warm-Cool** | **Luminance Entropy** | **Average Luminance** |
| Medial Left | $2.9056 \times 10^{-9}$ | 0.0592 | 0.1502 | 0.0588 | 0.0070 | 0.0335 | -0.0558 |

| | | | | | | | |
|---|---|---|---|---|---|---|---|
| Medial Right | $-2.7315 \times 10^{-9}$ | 0.0289 | 0.1976 | 0.0382 | -0.0094 | -0.0269 | 0.0213 |
| Lateral Left | $-1.2928 \times 10^{-8}$ | 0.0752 | 0.1574 | 0.0428 | 0.0043 | 0.0325 | -0.0601 |
| Lateral Right | $-8.6468 \times 10^{-9}$ | 0.0609 | 0.2223 | 0.0261 | 0.0152 | -0.0093 | -0.0156 |
| P7 | B0 | Average Saturation | Food | Circles | Warm-Cool | Luminance Entropy | Average Luminance |
| Medial Left | $1.2939 \times 10^{-8}$ | 0.0510 | 0.1334 | 0.0530 | 0.0308 | 0.0404 | -0.0551 |
| Medial Right | $5.8260 \times 10^{-10}$ | 0.0481 | 0.1942 | 0.0537 | 0.0244 | -0.0078 | -0.008 |
| Lateral Left | $8.1592 \times 10^{-9}$ | 0.0542 | 0.1524 | 0.0433 | 0.0094 | 0.0234 | -0.0227 |
| Lateral Right | $1.5644 \times 10^{-8}$ | 0.0346 | 0.2117 | 0.0244 | 0.0320 | -0.0063 | 0.0036 |
| P8 | B0 | Average Saturation | Food | Circles | Warm-Cool | Luminance Entropy | Average Luminance |
| Medial Left | $2.2001 \times 10^{-9}$ | 0.0577 | 0.1489 | 0.0452 | 0.0246 | 0.0696 | -0.0737 |
| Medial Right | $4.5941 \times 10^{-9}$ | 0.0390 | 0.1648 | 0.0321 | 0.0148 | 0.0057 | -0.0097 |
| Lateral Left | $5.0486 \times 10^{-9}$ | 0.0547 | 0.0986 | 0.0737 | 0.0204 | 0.0895 | -0.0916 |
| Lateral Right | $1.8202 \times 10^{-9}$ | 0.0426 | 0.1620 | 0.0251 | 0.0261 | 0.0161 | -0.0234 |

The beta coefficients for each multiple linear regression of the four streams for all eight participants.

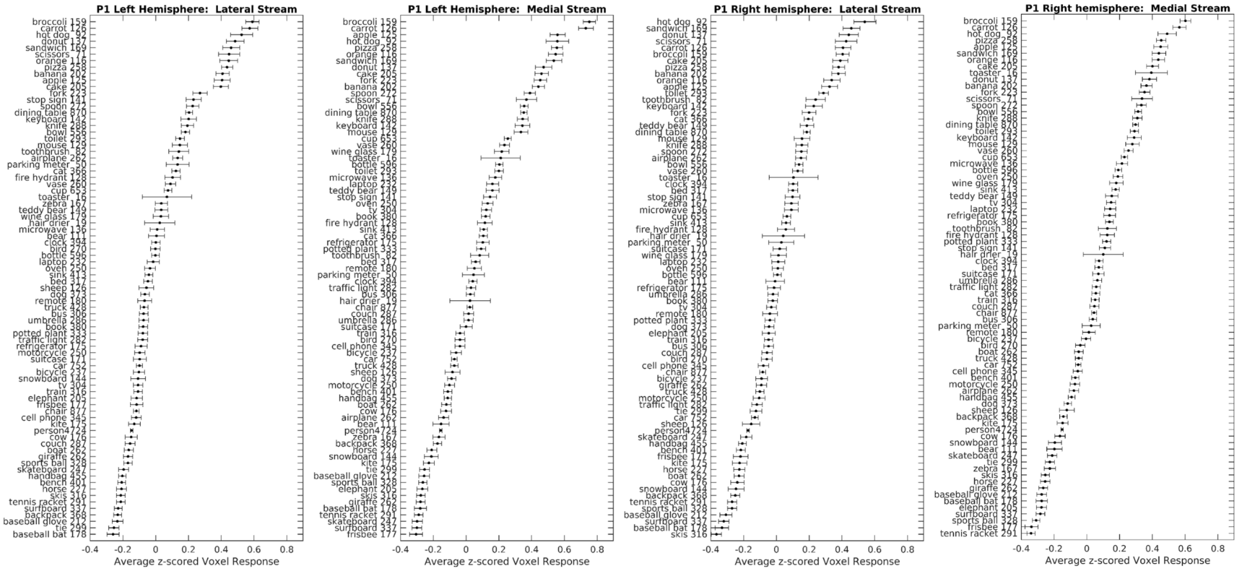
P1 Left Hemisphere: Lateral Stream
P1 Left Hemisphere: Medial Stream
P1 Right hemisphere: Lateral Stream
P1 Right hemisphere: Medial Stream
Average z-scored Voxel Response

65

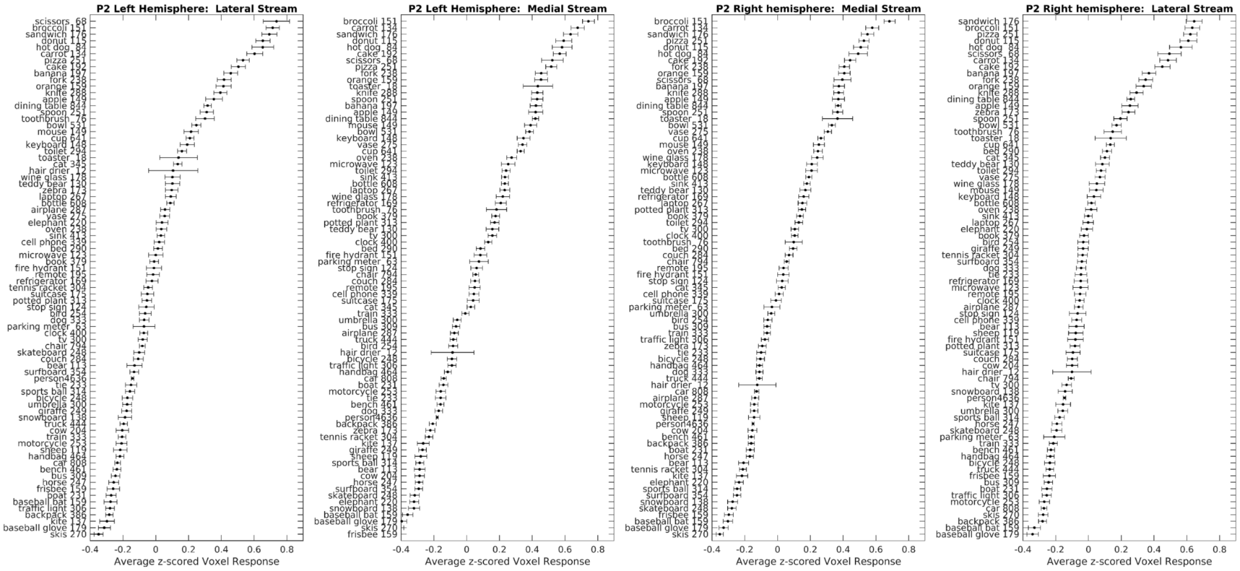
P2 Left Hemisphere: Lateral Stream
P2 Left Hemisphere: Medial Stream
P2 Right hemisphere: Medial Stream
P2 Right hemisphere: Lateral Stream
Average z-scored Voxel Response

66

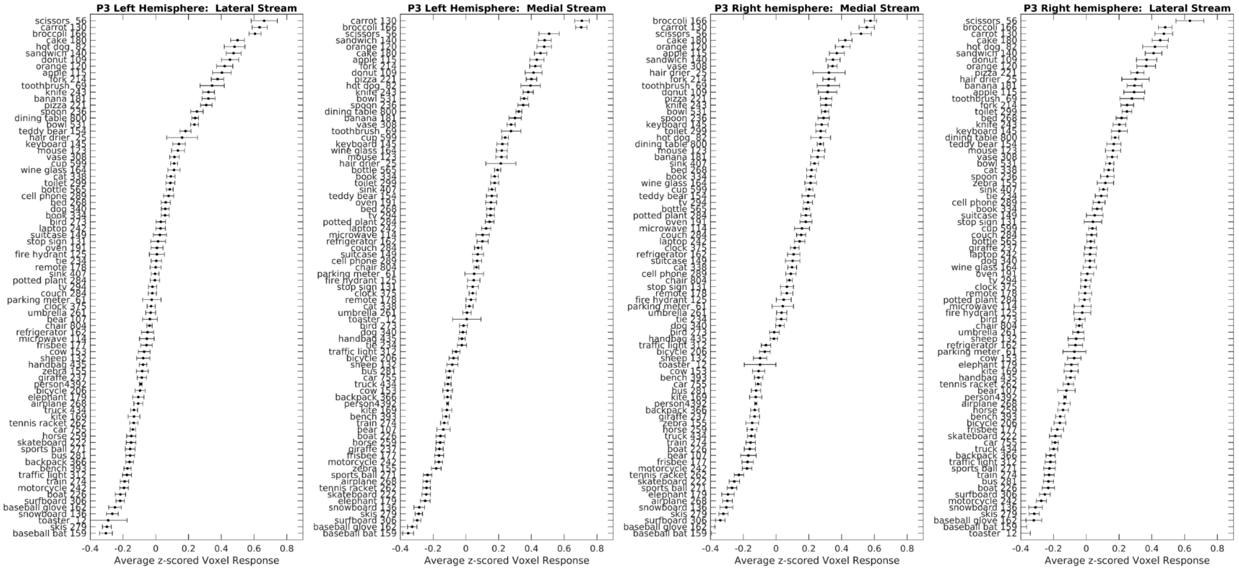
P3 Left Hemisphere: Lateral Stream
P3 Left Hemisphere: Medial Stream
P3 Right hemisphere: Medial Stream
P3 Right hemisphere: Lateral Stream
Average z-scored Voxel Response

67

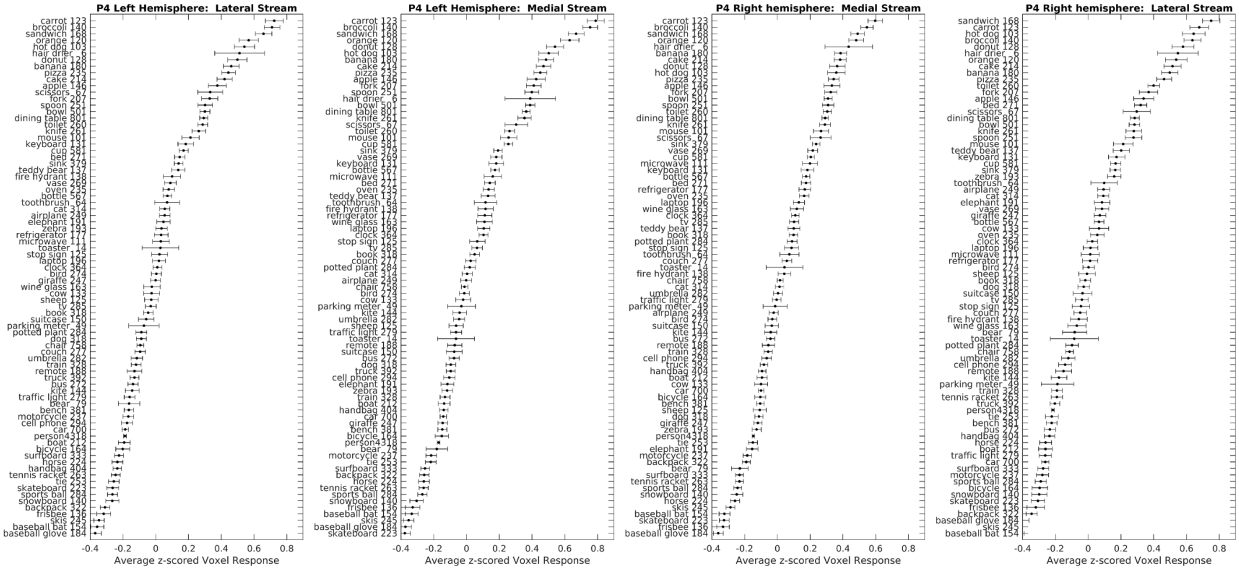
P4 Left Hemisphere: Lateral Stream
P4 Left Hemisphere: Medial Stream
P4 Right hemisphere: Medial Stream
P4 Right hemisphere: Lateral Stream
Average z-scored Voxel Response

68

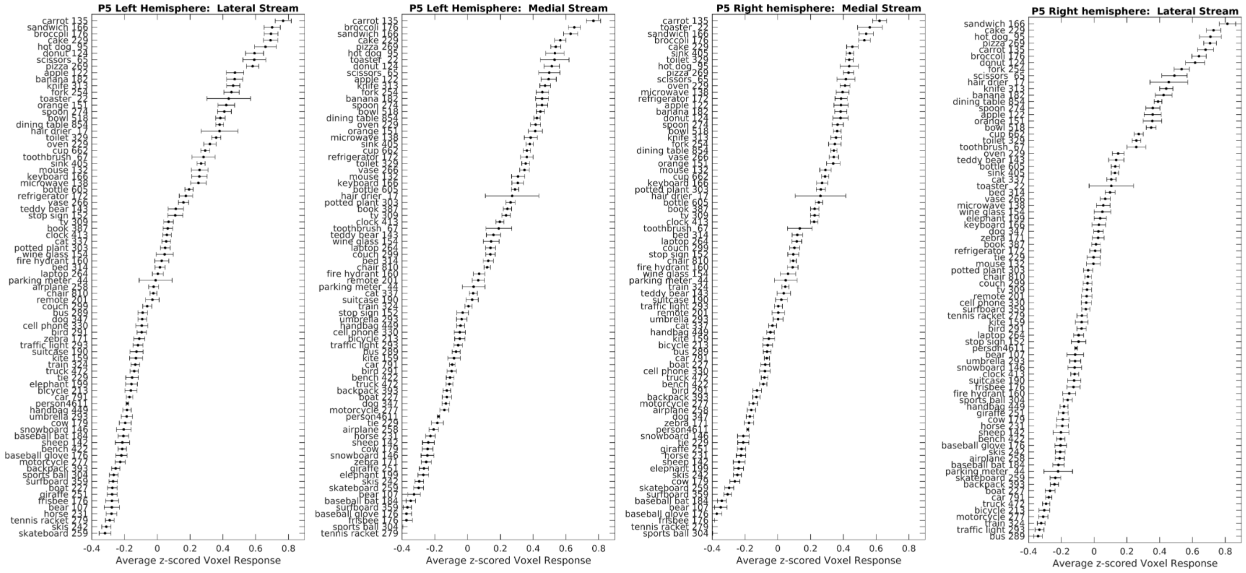
P5 Left Hemisphere: Lateral Stream
P5 Left Hemisphere: Medial Stream
P5 Right hemisphere: Medial Stream
P5 Right hemisphere: Lateral Stream
Average z-scored Voxel Response
Average z-scored Voxel Response
Average z-scored Voxel Response
Average z-scored Voxel Response

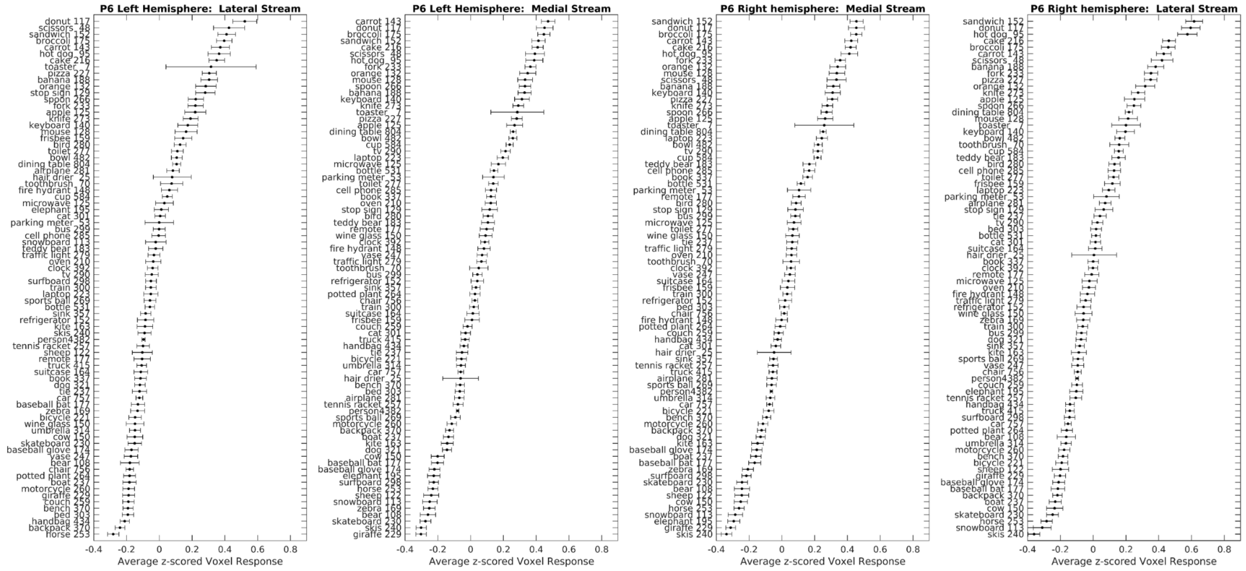
P6 Left Hemisphere: Lateral Stream
P6 Left Hemisphere: Medial Stream
P6 Right hemisphere: Medial Stream
P6 Right hemisphere: Lateral Stream
Average z-scored Voxel Response
Average z-scored Voxel Response
Average z-scored Voxel Response
Average z-scored Voxel Response

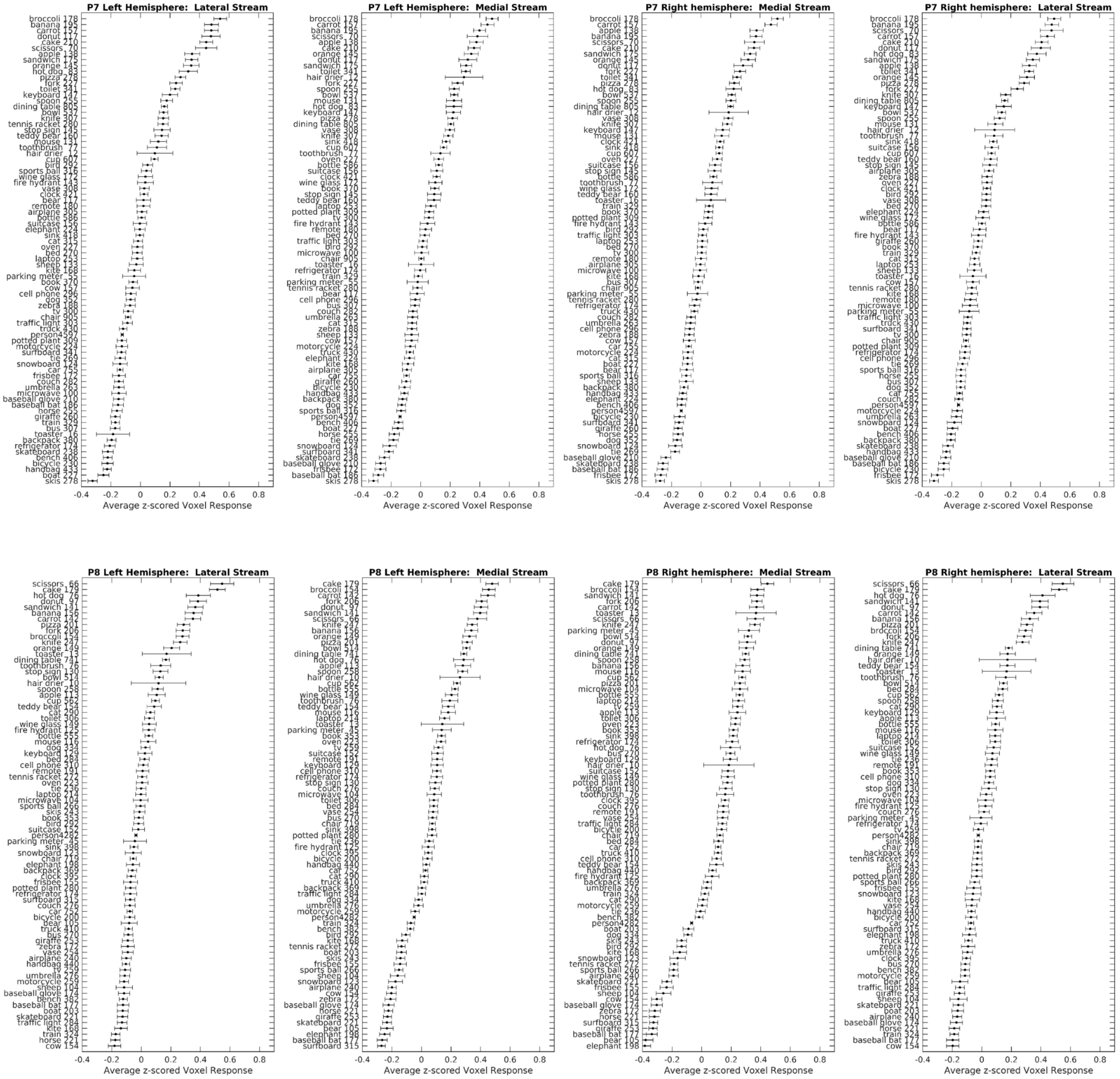


**Figure 7. Average voxel responses in the four ROIs for each COCO object category for each participant**

Each row of four panels shows average voxel response to each object category, with one panel for each ROI. Different rows are for different participants. The x-axis displays the average Z-scored voxel response, and the y-axis shows the object name and the number of images that contained this object for each participant. The 80 object categories are those identified and segmented in the COCO dataset. The object categories are ordered from those provoking the strongest response (top) to those provoking the weakest response (bottom). There may be multiple objects in one image. Error bars are standard errors of the mean.
